# Chromosome, Plasmid, or Both: Short-Term Dynamics and Long-Term Outcomes of Plasmid Cost Compensation under Trade-Offs

**DOI:** 10.64898/2026.07.06.736782

**Authors:** Christopher Witzany, Sebastian Bonhoeffer

## Abstract

Conjugative plasmids drive the dissemination of antimicrobial resistance genes and other conditionally beneficial accessory genes, but otherwise inflict fitness costs on their hosts that limit their spread. These costs can be ameliorated by compensatory mutations on the chromosome, the plasmid, or both combined (combined compensation). Mutants with plasmid-borne or chromosomal compensation differ in how they spread and compete, making the location of compensation an important determinant of plasmid and antimicrobial resistance (AMR) dynamics. We collate experimental data on plasmid-host co-evolution which, albeit limited, suggest compensatory benefits differ by location and are highest for combined compensation. We develop and analyse a mathematical model of compensatory evolution, finding that the long-term location of compensation is mainly determined by the highest cost reduction. Short term, however, succession dynamics arise from differences between locations: for instance, plasmid-borne compensation spreads horizontally, initially dominates, and can even facilitate the establishment of chromosomal or combined compensation. Strong trade-offs between compensation and either resistance or conjugation render compensation non-viable, but only conjugation trade-offs are location-dependent, disadvantaging plasmid-borne compensation and generating oscillatory dynamics. Our findings suggest plasmid-borne compensation may initially accelerate AMR-plasmid spread, whereas long-term chromosomal or combined compensation may enable hosts to accumulate multiple AMR-plasmids, promoting multidrug resistance.

## Introduction

The continuous spread and persistence of antimicrobial resistance genes (ARGs) poses a growing threat to global health [1–4]. A major driver of ARG spread, within and across bacterial communities, is horizontal gene transfer mediated by conjugative plasmids [5,6]. Moreover, conjugative plasmids increase the persistence of plasmid-borne ARGs in communities by transferring to plasmid-free cells, and can enable fixation of ARGs in the absence of antimicrobials [1,7,8].

Although plasmids often carry accessory genes that confer conditionally beneficial traits, such as antimicrobial resistance (AMR) or virulence factors, plasmids also inflict fitness costs on their bacterial hosts [9], for example due to increased translational or metabolic burden [10–12]. Plasmids can be lost stochastically during bacterial replication due to imperfect inheritance mechanisms, resulting occasionally in plasmid-free daughter cells. As plasmid carriage comes at a fitness cost, plasmid-free cells outcompete plasmid-carriers. Thus, for plasmid spread and persistence the conjugation rate needs to offset the plasmid loss and cost.

Empirical evidence shows that compensatory evolution ameliorates plasmid fitness costs [13,14], and thereby contributes to long-term persistence of plasmids [15]. Compensatory mutations can occur on the plasmid, the chromosome, or both [16–32]. The location of a compensatory mutation can have important consequences for plasmid spread and persistence in bacterial communities. Compensatory mutations located on the chromosome have been found to be not specific to only one type of plasmid, but rather lower the carriage costs of multiple different plasmids [23,25]. Thereby, a chromosomal compensatory mutation can enable or prolong the co-persistence of multiple plasmids within the same bacterial host cell (or host population). If these plasmids carry different ARGs chromosomal compensation can promote multidrug resistance [23] which in turn may lead to the emergence of multidrug resistance plasmids by within-host gene exchange [33]. Compensatory mutations located on the plasmid can reduce the plasmid’s cost across different bacterial strains and species, broadening its host-range [20,30]. Therefore, plasmid-borne compensatory mutations can facilitate the spread of ARGs to new, potentially pathogenic, bacterial hosts. Moreover, chromosomal and plasmid-borne compensation can emerge simultaneously within one host-plasmid population [16,23]. Combined compensation within the same host cell can lead to epistatic effects on plasmid costs ranging from synergism to antagonism. In particular it has been found that in some cases having compensation on both locations confers no additional benefit over having it at only one location [22,34]. Interactions between chromosomal and plasmid-borne mutations can thereby impact both plasmid host-range expansion and the host’s ability to tolerate plasmids, and thus shape plasmid success and evolution [16,35,36]. Understanding which factors determine the location of compensatory mutations is therefore relevant to understanding and predicting the spread and persistence of plasmids.

Experimental evolution studies find that both types of compensation occur, with chromosomal compensation being somewhat more frequent [13,18–21,29,30]. Theoretical studies, however, have argued that either strategy, plasmid-borne compensation or chromosomal compensation, could be the superior strategy. Zwanzig et al [37] argued that plasmid-borne compensation has the advantage that it can spread horizontally as well as vertically, whereas chromosomal mutations can only spread vertically. Others have argued [38–40] that chromosomal mutations should prevail, because hosts with a chromosomal mutation can transfer the plasmid to hosts lacking compensatory mutations, thus inflicting a fitness cost and increasing the competitive advantage of the chromosomally compensated hosts.

The biological reality may be even more complex than described in these theoretical studies, because compensatory mutations can reduce plasmid costs by reducing plasmid-encoded functions, such as conjugation or antibiotic resistance, thereby creating potential trade-offs between improved growth and these functions [11,27,41,42]. If such trade-offs depend on the location of the compensatory mutation, they may shape which location is favoured. For example, it is conceivable that mutations on the plasmid are more likely to directly affect plasmid-encoded traits such as conjugation or antibiotic resistance than mutations on the chromosome, which could make plasmid-borne compensation less favourable.

Here, we develop a mathematical model to investigate the competitive dynamics of compensatory mutations emerging on the plasmid, chromosome or both, and the factors that ultimately determine where compensation fixates. We parametrize our model by collating empirical values of plasmid fitness costs and cost amelioration associated with compensatory mutations from the literature. Our simulations show that the main determinant of where a compensatory mutation ultimately fixes is where compensation can reduce carriage cost most effectively. However, the dynamics can be complex: plasmid-borne compensation generally establishes faster but in the long term can be replaced by or combined with chromosomal compensation if this confers greater fitness gains. Trade-offs between compensation and conjugation can further complicate successions and even generate oscillatory dynamics, whereas trade-offs between compensation and resistance generally simplify successions because antibiotics remove plasmid-free cells and thereby reduce the importance of conjugation. Overall, our results suggest that the carriage cost reduction is the key determinant of how compensatory mutations shape plasmid spread and persistence, while also providing broader insight into the dynamics of two-host, two-plasmid systems and plasmid-host coevolution.

## Methods

### Literature review of plasmid costs and benefits of compensatory mutations

We reviewed experimental evolution studies investigating plasmid-host co-evolution. We identified 13 studies from which we could extract proxy fitness measures quantified by growth rate (CFU or OD data) or competition assays between combinations of ancestral and evolved and plasmid-free and plasmid-carrying bacteria (Supplemental Data File 1). These fitness measures (*λ*_*i*_) included ratios of Malthusian parameters [43] from single-strain [26,27] or competition experiments [19–21,24,29,32,44–47], and in one case a ratio of areas under the curve (AUC) of growth curves [48]. Where available, we treated individual evolved lines as separate measurements to capture within-experiment variance and compared this with averaging measurements per study.

Uncompensated plasmid fitness costs 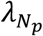 were determined by the growth rate of plasmid-carrying ancestor, i.e., before the evolution experiment, relative to the plasmid-free ancestor. Location of the compensatory mutations was deduced from the experimental observations reported by the studies as follows: If the evolved plasmid showed reduced costs in the ancestral host relative to the ancestral plasmid, it was attributed to plasmid-borne compensation 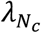. Vice versa if the ancestral plasmid showed reduced cost in the evolved host, we classified that as chromosomal compensation 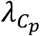. The effect of combined compensation (i.e., both compensations occurring simultaneously, 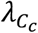) was assessed by comparing the evolved plasmid-host pair with the ancestral-evolved pairs. All fitness measures were normalized to that of the plasmid-free ancestor (i.e., 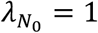). Lastly, the effect of chromosomal compensation in the absence of any plasmid 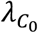 was set by the evolved and plasmid-cured strain, if reported. However, not all studies quantified fitness for all possible plasmid-host combinations. Note that the number of generations between the experiments, selection regimes, population sizes, bacterial host species and plasmids differ.

### Population model

To investigate the factors that determine whether a compensatory mutation should establish on the plasmid, on the chromosome, or both – and how fast – we built a mathematical population model that allows for these mutations to emerge, compete and interact with each other. See Figure 1 for a visual overview of the model and Text S1 for equations. We model two types of bacterial hosts: one ancestral (*N*) and one with a compensatory mutation on the chromosome (*C*). Both host types can carry either the ancestral plasmid (*N*_*p*_, *C*_*p*_), a plasmid with a compensatory mutation (*N*_*c*_, *C*_*c*_), or no plasmid (*N*_0_, *C*_0_).

**Figure 1.**
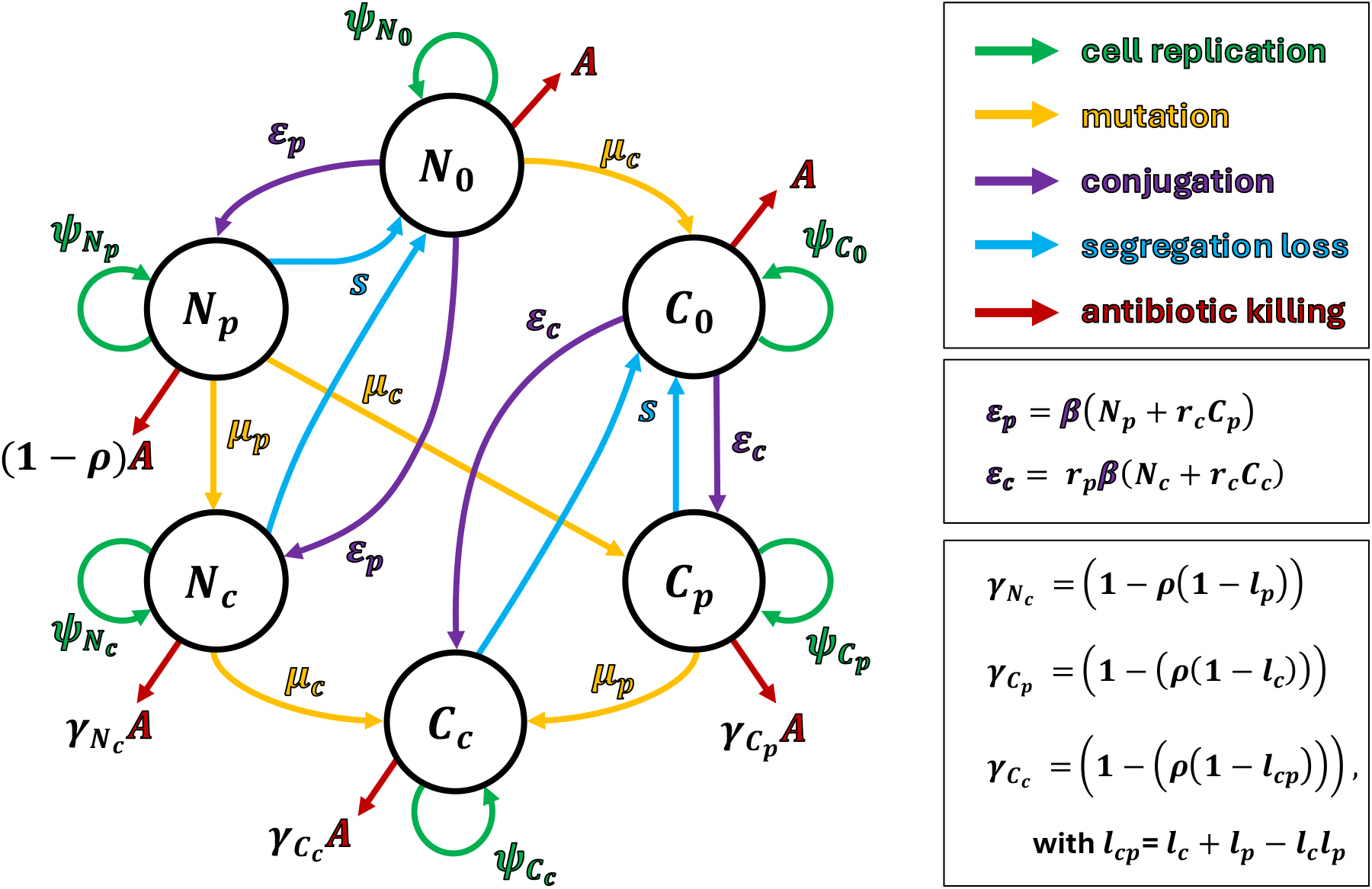
Diagram of the full model of plasmid cost compensatory evolution in different locations. The model describes the dynamics of six bacterial populations *N*_0_, *N*_*p*_, *N*_*c*_, *C*_0_, *C*_*p*_ and *C*_*c*_. Cells with the ancestral chromosome are denoted by “N” and those with a chromosomal compensatory mutation located on the chromosome by “C”. Plasmid carriage is indicated by subscripts: “0” (no carriage), “p” (ancestral plasmid), and “c” (compensated plasmid). Processes are grouped by classes and color-coded: cell replication, (compensatory) mutation, plasmid conjugation, segregation loss, antibiotic killing (including plasmid-conferred resistance). See Methods for a detailed description of the model, Text S1 for model equations, and Table S1 for an overview of parameter definitions and values.

Each of the resulting six populations has a unique growth rate *λ*_*i*_ relative to that of the plasmid-free ancestor 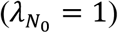. The uncompensated plasmid-carrying population *N*_*p*_ has growth rate 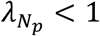, reflecting the plasmid carriage cost. The fitness effects of mutations on the chromosome, plasmid, or both are captured by the growth rates of the corresponding populations, i.e., 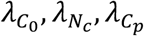, and 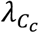. Whether a mutation compensates for plasmid cost is determined by whether the corresponding populations growth rate exceeds 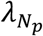, and for chromosomal mutations 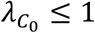 must additionally hold, i.e., chromosomal mutations are beneficially only under plasmid carriage [49].

All plasmid-bearing populations can lose the plasmid via growth dependent segregations loss (*s*). Compensation on the plasmid (*C*_*c*_ or *N*_*c*_) can emerge from plasmid-bearing populations (*N*_*p*_ and *C*_*p*_) via growth dependent mutation by rate *μ*_*p*_. Analogously chromosomal compensation can emerge by rate *μ*_*c*_ from ancestral host populations *N*_0_, *N*_*p*_, and *N*_*c*_.

We model selection pressure exerted by antibiotics as a killing rate *A*. The plasmid carries a resistance gene, which reduces the killing rate to (1 − *ρ*)*A*, with *ρ* = 0 reflecting no resistance and *ρ* = 1 complete resistance. Since resistance genes can contribute to plasmid costs, compensatory mutations can mitigate these costs by reducing expression of ARGs. Therefore, we model that plasmid compensation (*N*_*c*_) or chromosomal compensation (*C*_*p*_) can result in reduced resistance, described by a proportional reduction of *ρ* with magnitude *l*_*c*_ or *l*_*p*_, respectively, where *l*_*c*_ and *l*_*p*_ can range from 0 (no reduction) to 1 (complete loss of resistance). Having combined compensation (*C*_*c*_) results in a reduction of resistance described by *l*_*c*_ + *l*_*p*_ − *l*_*c*_*l*_*p*_, a simple mathematical formulation chosen here to ensures that combined loss of resistance is greater than each individual loss but does not exceed 1. Taken together, the effective antibiotic killing rate for any population can therefore be described by (1 − *ρ*(1 − *l*_*x*_))*A*.

Plasmids can be transmitted horizontally from plasmid-bearing cells (*N*_*p*_, *N*_*c*_, *C*_*p*_, and *C*_*c*_) to plasmid-free cells (*N*_0_ and *C*_0_) by conjugation at rate β following mass-action dynamics. Since conjugation is a costly process, compensatory mutations on the chromosome or plasmid can reduce costs by reducing conjugation. Here we model this by decreasing the conjugation rate by factors *r*_*c*_ for chromosomal mutations and *r*_*p*_ for plasmid mutations, respectively, ranging from 0 (full loss of conjugation) to 1 (no change in conjugation). The conjugation rate of *C*_*c*_ is described by *r*_*p*_*r*_*c*_*β*.

We introduce a density-dependent death rate *ϕ*, that equals the net growth in overall population size thereby enforcing competition by limiting the overall population size to 1. Thus, the population sizes of the individual subpopulations represent proportions of the overall population. Our model therefore mimics the conditions of a turbidostat by maintaining a constant population size through proportional washout.

### Analytical Solutions

First, to make the model analytically tractable, we implemented a simplified version in *Mathematica* (Version 13.0.1.0) that neglects mutations, segregation loss, and changes in antibiotic resistance or conjugation rates associated with compensatory mutations (i.e., *μ*_*c*_, *μ*_*p*_, *l*_*c*_, l_*p*_, *s* = 0 and *r*_*c*_, *r*_*p*_ = 1). Further, we assume that all plasmid carrying populations have full resistance (*ρ* = 1). For this simplified model, we determined all biologically feasible steady states (i.e., steady states based on biologically meaningful parameters and non-negative population sizes). Then, we conducted an invasion analysis to determine the parameter conditions under which any of the six populations can successfully invade a given steady state. Briefly, a population can invade, and the steady state is unstable, if the per-capita growth rate of the invading population is positive when all other populations are at their corresponding steady state frequency.

### Numerical Simulations

Next, we implemented the full model in *Julia* (v. 1.11.5) using *DifferentialEquations*.*jl* (v. 7.16.1) [50]. Time was measured in hours; parameters are specified in Table S1. We ran numerical simulations starting from ancestral populations of plasmid-free and plasmid-bearing bacteria (*N*_0_ = 0.75, *N*_*p*_ = 0.25, *N*_*c*_ = *C*_0_ = *C*_*p*_ = *C*_*c*_ = 0) and run simulations up to t=10,000h unless otherwise indicated. We categorize simulation results by 1) plasmid persistence, 2) dominant population, and 3) the succession of dominant plasmid-carrying populations. Plasmid persistence or loss is determined by whether the sum of plasmid-carrying populations (*N*_*p*_, *N*_*c*_, *C*_*p*_, *C*_*c*_) at the end of the simulation exceeded 0.1% in frequency. The dominant population is defined as the largest population at the end of simulation. The succession of dominant plasmid-carrying populations is contingent on the plasmid persistence and is the sequence of the largest plasmid-carrying populations over time, thereby describing which plasmid population succeeds which. We conducted several sensitivity analyses to assess how sensitive our numerical results are to parameter values (see Supplementary Text S2 for details).

### Trade-off between compensatory benefits and resistance or conjugation

Mutations that reduce – i.e. compensate – plasmid carriage costs may do so by reducing metabolically expensive antibiotic resistance or plasmid conjugation (Figure S1). We implement such trade-offs by assuming that a given compensatory benefit (Δ_*c*_) is due to either a loss of resistance (*l*_*c*_, *l*_*p*_) or, alternatively, a reduction of conjugation rate (*r*_*c*_, *r*_*p*_) following a simple power-law. Specifically, for the resistance trade-off, we set 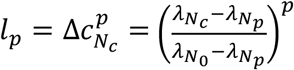 and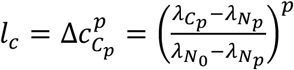, corresponding to mutations located on the plasmid or the chromosome, respectively. The exponent *p* controls the trade-off strength, with values of *p* > 1 resulting in weak (concave) and *p* < 1 in strong (convex) trade-off curves (Figure S1). When mutations yield growth rates below 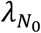 and above 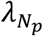, *l*_*p*_ and *l*_*c*_ are < 1. However, this condition is not always met in the empirical data (Figure 2). Hence, for simplicity, mutations yielding growth rates exceeding 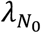 or falling below 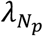 are assigned Δ_*c*_ = 1 or Δ_*c*_ = 0, respectively, thereby preventing increases in resistance or conjugation relative to *N*_*p*_. Note, that for *C*_*c*_ loss of resistance or conjugation emerges from the combined effects of the single mutations (see descriptions above, for equations Text S1). The conjugation trade-off is analogously implemented with 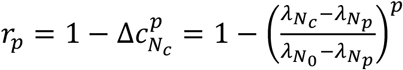 and 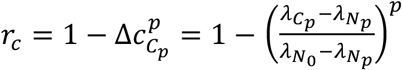.

**Figure 2.**
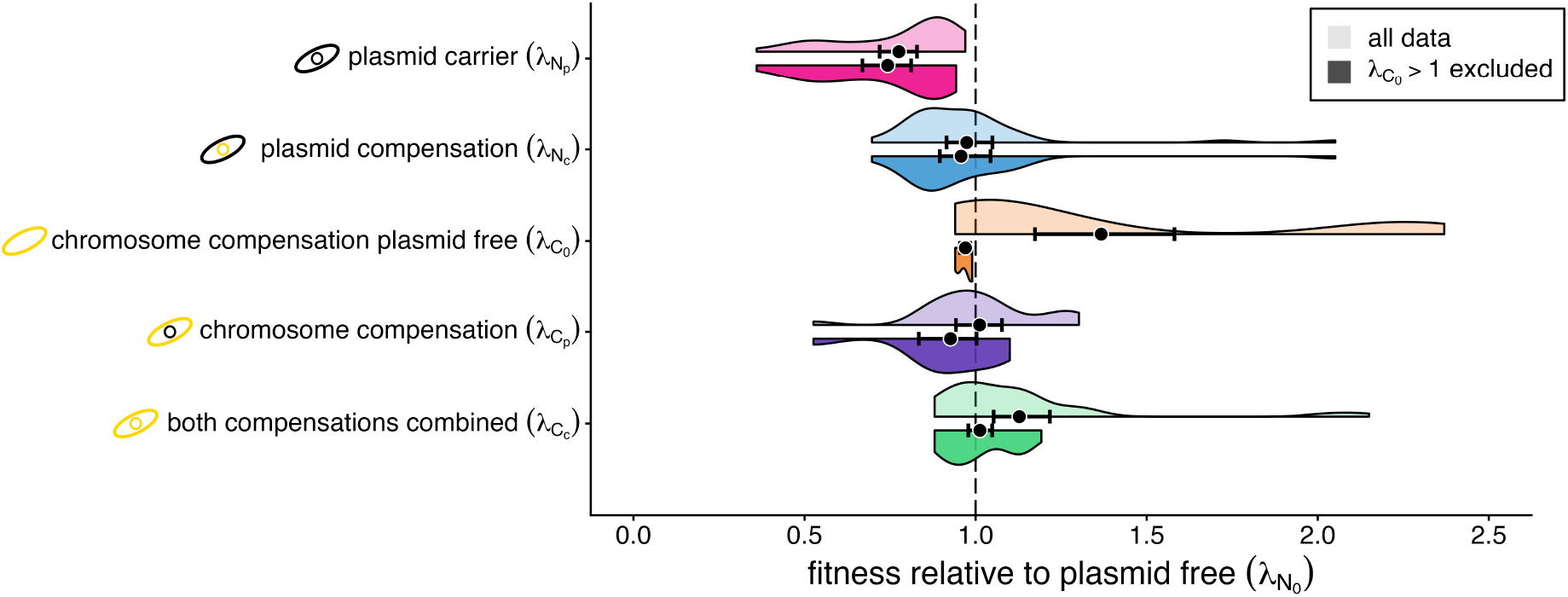
Fitness costs and amelioration of plasmid carriage before and after experimental evolution. Composite kernel density (violin) plots showing fitness measures, compiled from the literature, normalized to the respective plasmid free ancestral bacterial host (*N*_0_). Shown are fitness costs of plasmid carriage before experimental evolution (*N*_*p*_) and after experimental evolution, grouped by the reported genetic location of compensatory evolution, i.e., on the plasmid (*N*_*c*_), on the chromosome (*C*_*p*_), or on both the chromosome and the plasmid simultaneously (*C*_*c*_). Note that where the location is not specified or inferable from the original studies, the location is conservatively attributed to *C*_*c*_. *C*_0_ represents the fitness effects of chromosomal compensatory mutations in the absence of any plasmid, i.e., the fitness of the evolved host after being cured of the plasmid. Upper, lighter-coloured half-violin plots show all data points, whereas the lower, darker-coloured half-violins exclude cases where chromosomal compensatory mutations result in fitness greater than the plasmid free ancestor (*C*_0_ > 1). Mean fitness values for each group (black points) are shown with bootstrapped 95% confidence intervals. All violin plots are scaled to the same maximum width. See Supplemental Data File 1 for underlying data and Methods for details on literature review and data extraction.

## Code Availability

All code used for simulations and analyses is available at https://github.com/ChrisWitzany/chrom_plasm_or_both.

## Results

### Empirical measures suggest that highest fitness is conferred if compensatory mutations occur on both, chromosome and plasmid

To quantify costs and benefits of plasmid carriage, we compiled fitness measures before and after experimental evolution from 13 published studies [19–21,24,26,27,29,32,44–48] (Supplemental Data File 1). We find that, before experimental evolution, the mean fitness of plasmid carrying bacteria (*N*_*p*_) relative to the plasmid free ancestor (*N*_0_) is 0.78 (Figure 2, upper panel, lighter half-violin plots). After evolution, *C*_*c*_ (compensation on both the plasmid and chromosome) confers the highest average compensation of plasmid costs, with a mean fitness of 1.13, while *N*_*c*_ (compensation on plasmid) and *C*_*p*_ (compensation on the chromosome) have a mean fitness of 0.97 and 1.01, respectively. However, *C*_0_, i.e., the evolved host cured of the evolved plasmid, has overall the highest mean fitness of 1.37. This suggests that, at least in part, general beneficial chromosomal mutations result in the increased fitness of both *C*_0_ and *C*_*c*_ rather than mutations that specifically compensate plasmid costs. This conflicts with the strict definition of a compensatory mutations, which is that that compensatory mutations are deleterious or at most neutral in the absence of what they compensate (i.e., there is positive epistasis) [49]. In other words, a true compensatory mutation located on the chromosome is expected to be deleterious, or at most neutral, in absence of the plasmid (i.e., 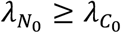). Excluding the sets of fitness measurements where *C*_0_ > 1 reduces the mean fitness of all groups, but substantially only for *C*_0_, which drops to 0.97 (Figure 2, lower, darker half-violin plots). Despite this reduction, the highest mean fitness is still conferred by combined chromosomal and plasmid compensations (*C*_*c*_, 1.01), followed by plasmid compensations (*N*_*c*_, 0.96), and chromosomal compensations (*C*_*p*_, 0.93) (Table S2). Restricting this analysis to only studies reporting head-to-head competition fitness measures does not change these findings qualitatively (Figure S2, Table S3), and neither does averaging measures by study (Figure S3, Table S4).

### Compensatory mutations are invadable except where benefits are highest

To understand whether plasmid costs and compensatory fitness effects alone determine where compensatory mutations are expected to be located long term – on the plasmid, the chromosome, or on both – we first conduct an analytical invasion analysis. For this, we neglect segregation loss and mutation rates (*μ*_*c*_, *μ*_*p*_, *l*_*c*_, *l*_*p*_, *s* = 0) and assume no trade-offs (*l*_*c*_, *l*_*p*_, *s* = 0, *r*_*c*_, *r*_*p*_ = 1). We find six steady states with only a single population present (*N*_0_, *N*_*p*_, *N*_*c*_, *C*_0_, *C*_*p*_, *C*_*c*_, Figure 3A) and four steady states with three populations coexisting (*N*_0_, *N*_*c*_, *C*_*c*_; *N*_0_, *N*_*p*_, *C*_*p*_; *N*_*c*_, *C*_0_, *C*_*c*_; and *N*_*p*_, *C*_0_, *C*_*p*_, Figure 3B,C). Interestingly, compensated and non-compensated chromosomes can coexist, while compensated and non-compensated plasmids cannot (see Text S3).

**Figure 3.**
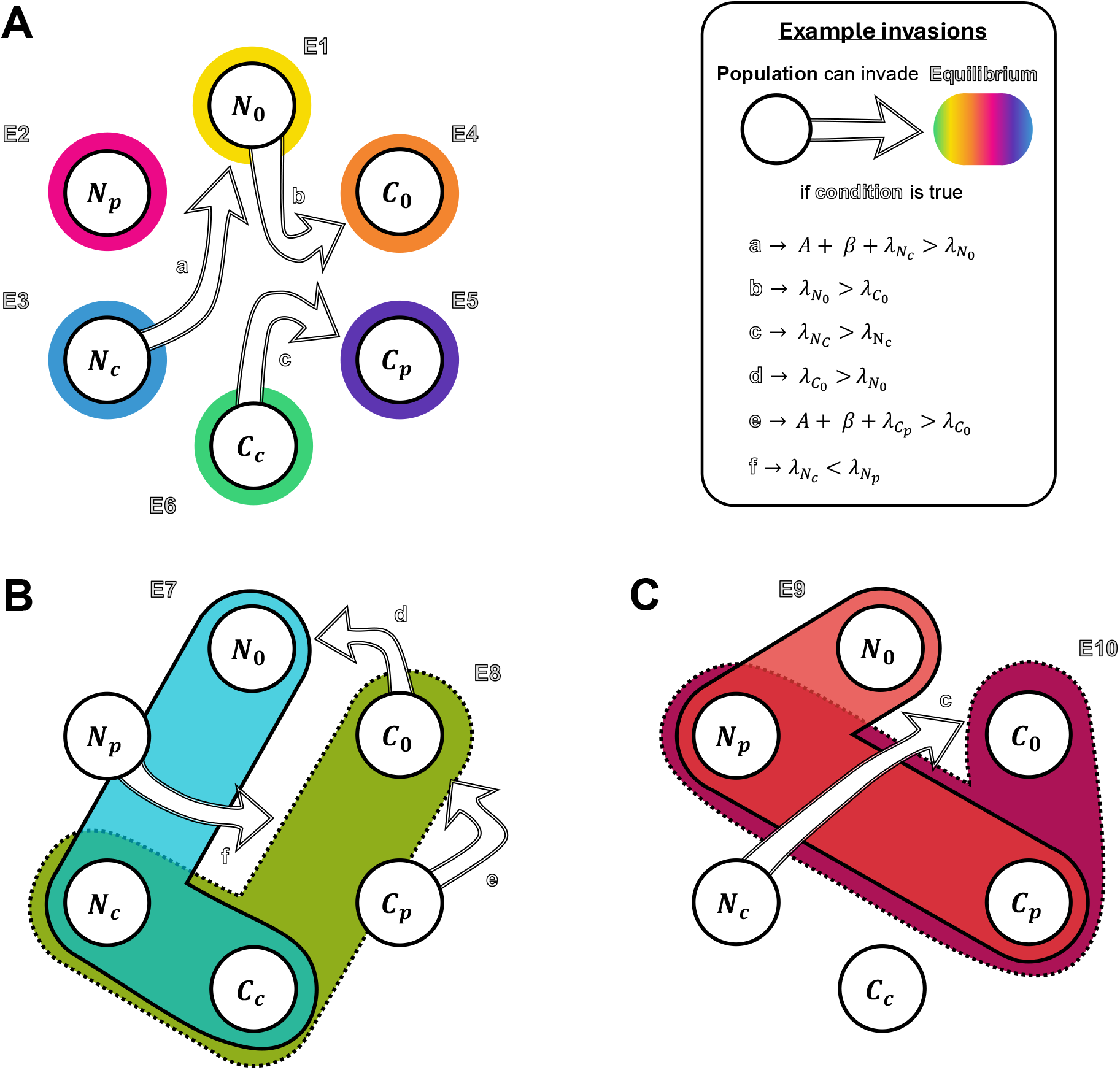
Biologically feasible equilibria and example invasion criteria of the simplified model. Each of the six populations, *N*_0_, *C*_0_, *C*_*p*_, *C*_*c*_, *N*_*c*_, and *N*_*p*_, is represented by a circle (see Figure 1 for details on the populations). The 10 possible equilibria, E1 to E10, are shown by semi-transparent coloured outlines. Populations enclosed within the same outline are present at the corresponding steady state. The six single population equilibria (E1 to E6) are shown in (**A)**, and the three-population equilibria corresponding to both compensations (E7 and E8) and chromosome compensation only (E9 and E10) in **B** and **C**, respectively. Invasion of a single population into a steady state, shown by arrows originating from the invader population and pointing to the steady state, is possible if the corresponding parameter condition (a-f) is met. *A* represents antibiotic killing, *β* the conjugation rate, and *λ* the respective population-specific growth rate; see Table S1 for descriptions of all parameters. Note that only a subset of representative invasion criteria is shown. See Table S2 for a full list of all invasion criteria.

Overall, there is no steady state that cannot be invaded. Regardless of where compensation occurs, the invasion analysis of the steady states shows that plasmid-carrying populations with compensatory mutations (*N*_*c*_, *C*_*p*_, or *C*_*c*_) can invade the ancestral plasmid (*N*_*p*_) only if they have a higher growth rate (*λ*_*i*_). Likewise, *N*_*c*_, *C*_*p*_, and *C*_*c*_ can invade each other only if they have a higher growth rate (*λ*_*i*_). More generally, the plasmid-free ancestor (*N*_0_) can be invaded by plasmid-carrying populations if antibiotic selection (A) is strong enough, conjugation (β) is sufficiently high, and the invader has a sufficiently high growth rate (*λ*_*i*_) relative to *N*_0_, i.e., 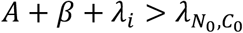, where 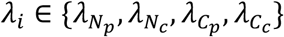. The three-population steady states can be invaded by any absent population if the invader outcompetes one of the three resident populations. This results in multiple invasion routes (see Figure 3B,C for examples), suggesting that co-existence of compensated and uncompensated populations is easily destabilized. See Supplementary Text S3 for a detailed description of the invasion criteria and analysis. Taken together, as no steady state is universally immune to invasion, no compensation location is a priori superior, suggesting that population dynamics could play an important role in compensatory evolution – which we investigate next.

### Plasmid-borne compensation establishes faster than compensation on the chromosome and wins under equal benefits

To investigate the temporal dynamics of compensatory evolution in detail, we relax our simplifying assumptions that segregation loss (*s*) and mutation rates (*μ*_*p*_, *μ*_*c*_) are negligible and turn to numerical simulations. We start all our simulations from initial conditions where ancestral plasmids are present at minority frequency (*N*_0_ = 0.75, *N*_*p*_ = 0.25, and all other populations are equal to zero). We consider a costly conjugative plasmid that cannot persist without compensatory evolution (*μ*_*c*_ = *μ*_*p*_ = 0): *N*_*p*_ declines and eventually goes extinct (Figure 4A) because conjugation cannot offset plasmid loss and carriage cost.

**Figure 4.**
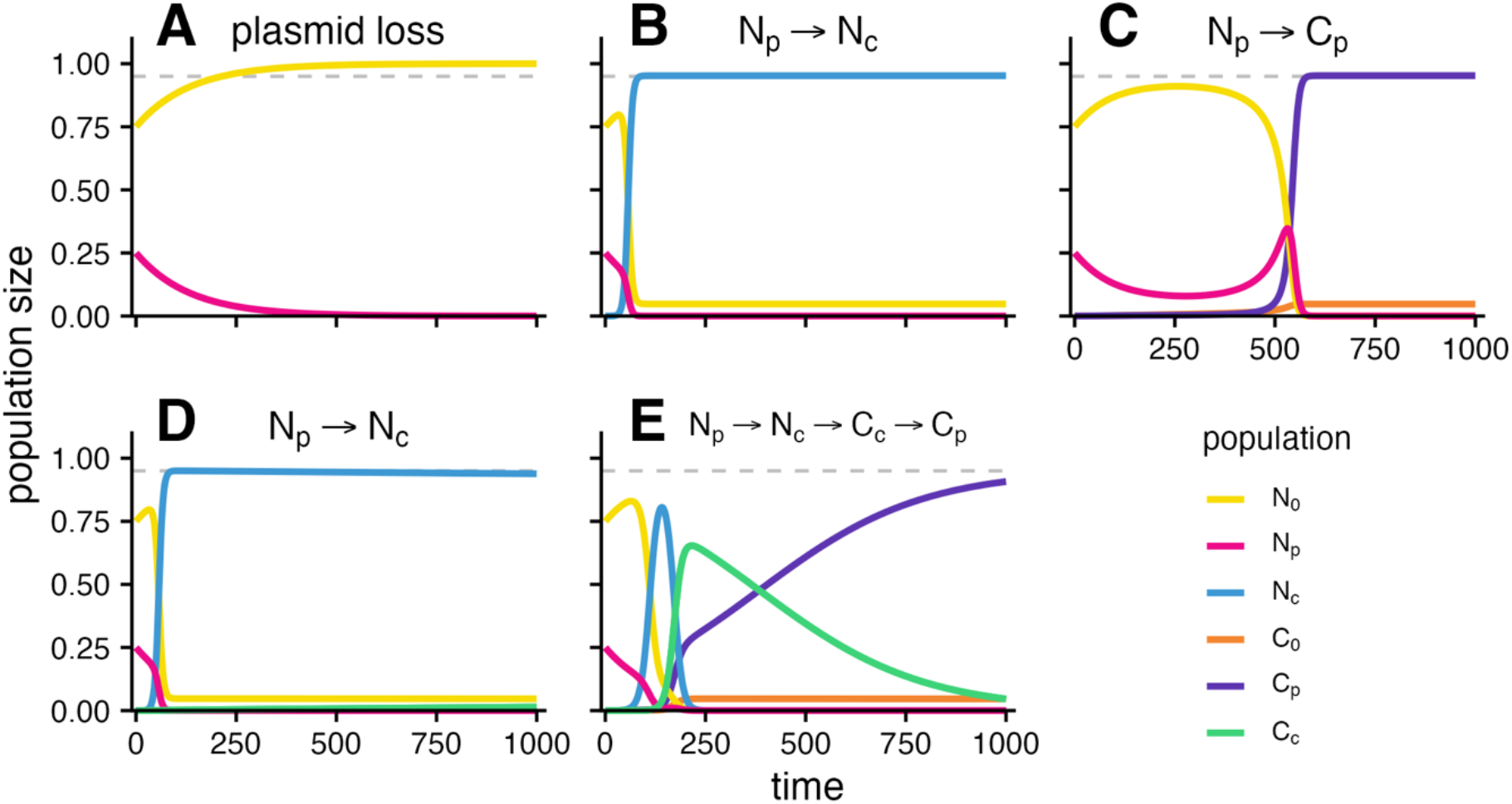
Temporal dynamics of compensatory evolution for beneficial mutations emerging on the plasmid and chromosome. The population sizes over time (h) are shown for the following cell types: ancestral, plasmid-free cells (*N*_0_, yellow), cells carrying the uncompensated plasmid (*N*_*p*_, pink), cells carrying the compensated plasmid (*N*_*c*_, blue), plasmid-free cells with a compensatory mutation on the chromosome (*C*_0_, orange), and cells carrying the compensated plasmid and have a compensatory mutation on the chromosome (*C*_*c*_, green). Panels A-E correspond to different scenarios reflecting whether compensatory mutations are possible, and if so, where. (**A**) Emergence of compensatory mutations is not possible, i.e., both mutation rates are zero (*μ*_*c*_ = *μ*_*p*_ = 0). (**B**) Only compensatory mutation on the plasmid is possible (*μ*_*c*_ = 0, *μ*_*p*_ = 1.5 × 10^−5^). (**C**) Only compensatory mutation on the chromosome is possible (*μ*_*c*_ = 1.5 × 10^−5^, *μ*_*p*_ = 0). (**D-E**) Both compensatory mutations are possible (*μ*_*c*_ = 1.5 × 10^−5^, *μ*_*p*_ = 1.5 × 10^−5^) with identical (**D**) or differing benefits (**E**). In A-D the benefits of the compensatory mutations in each location – i.e., the increase in growth rate relative to *N*_*p*_ – are identical 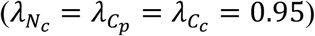. In **E** chromosomal compensation confers the highest benefit and plasmid-born the lowest 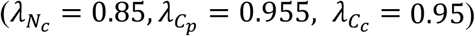. The grey, dashed, horizontal line indicates a population size of 0.95. All simulations start with *N*_0_ = 0.75, *N*_*p*_ = 0.25, *N*_*c*_ = *C*_0_ = *C*_*p*_ = *C*_*c*_ = 0. See Methods for a description of the model and see Table S1 for all parameters used.

If a compensatory mutation can emerge only on the plasmid (*μ*_*c*_ = 0, *μ*_*p*_ = 10^−5^), *N*_*c*_ quickly emerges and reaches over 95% prevalence, causing a rapid decline of *N*_*p*_ as well as *N*_0_, which does not go extinct however (Figure 4B). If instead only an equally beneficial compensatory mutation on the chromosome is possible (*μ*_*c*_ = 10^−5^, *μ*_*p*_ = 0), *C*_*p*_ emerges much later and reaches 95% prevalence only after a substantial delay (about 6.5-fold slower, Figure 4C). This is because *N*_*c*_ can spread both vertically through replication and horizontally through conjugation with *N*_0_, whereas *C*_*p*_ can only spread vertically – because conjugation between *N*_0_ and *C*_*p*_ results in *N*_*p*_, not *C*_*p*_. Conjugation between *C*_*p*_ and *N*_0_ does, however, give *C*_*p*_ an indirect advantage by converting fitter *N*_0_ into less fit *N*_*p*_, which *C*_*p*_ can then outcompete since 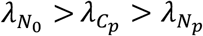 (Figure 3C; sometimes called plasmid “weaponisation”[40,51]). Consequently, conjugation between *C*_*p*_ and *N*_0_ initially slows and then reverses the decline of *N*_*p*_, resulting in a peak in population size of 0.35 for the given example, that coincides with the increase of *C*_*p*_, but shortly after this peak, both *N*_*p*_ and *N*_0_ ultimately go extinct. This advantage is lessened by increasing segregation rate *s*, as *N*_*p*_ loses the plasmid faster and reverts more quickly to *N*_0_ (Figure S4). Moreover, while both *N*_*c*_ and *C*_*p*_ experience segregation loss and continuously generate plasmid-free cells (*C*_0_ from *C*_*p*_ and *N*_0_ from *N*_*c*_, Figure 4B,C), this is more detrimental to *C*_*p*_ (Figure S4), because chromosomal compensation is costly in absence of plasmids (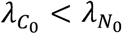, Figure S5).

If compensatory mutations can emerge on the plasmid and the chromosome (*μ*_*c*_ = 10^−5^, *μ*_*p*_ = 10^−5^), *N*_*c*_ quickly establishes (Figure 4D) – as expected from the case when only plasmid compensation is possible (Figure 4B). Despite *C*_*p*_ and *C*_*c*_ conferring equal benefits to *N*_*c*_, *C*_*p*_ does not increase and *C*_*c*_ only increases extremely slowly by continuous mutation of *N*_*c*_ to *C*_*c*_. Over very long times scales (up to 10^6^ h), whether *C*_*c*_ overtakes *N*_*c*_ or remains at a lower frequency depends on whether 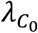 and *s* are sufficiently small (Figure S6), as *C*_*c*_ is constrained by the cost of chromosomal compensation (same as *C*_*p*_, see above). If benefits differ, complex dynamics can emerge, with multiple transiently dominant populations (Figure 4E). Interestingly, early spread of *N*_*c*_ can even facilitate and speed up the later establishment of *C*_*p*_ (Figure S7). This may seem counterintuitive, but it is related to the indirect advantage of *C*_*p*_ in isolation described above (plasmid “weaponisation”). There, conjugation converts fitter *N*_0_ into less fit *N*_*p*_. In this case, however, conjugation between *N*_*c*_ and *N*_0_ cells yield additional *N*_*c*_ cells, which can then themselves conjugate with *N*_0_ and generate even more *N*_*c*_. This self-amplification makes the effect much stronger than the indirect advantage of *C*_*p*_ in isolation; thus *N*_*c*_ rapidly depletes *N*_0_. Once *N*_0_ cells are depleted, *C*_*p*_ can in turn replace *N*_*c*_ if it has a higher growth rate, which can be faster than displacing *N*_0_ depending on the relative growth rate hierarchy (Figure S8). Overall, if compensatory benefits are equal, plasmid-borne compensation is more likely to establish and does so faster than chromosomal compensation across a wide range of parameters (Figure S9).

### Plasmid-borne compensation typically drives initial spread but can be succeeded by chromosomal or combined compensation

So far, our numerical simulations have focused on equal benefits across locations (with the exception of Figure 4E). However, empirical data suggest that benefits can differ by location (Figure 2). Therefore, we next investigate in detail how differences in benefits shape which compensatory mutations increase and spread. For this, we start with the same conjugative plasmid that cannot persist without compensation from Figure 4A (Table S1) and run simulations across combinations of chromosomal 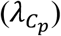 and plasmid compensatory benefits 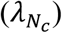 – each ranging from 0.7 to 1 (range informed by empirical data Figure 2; see Figure 5 for simulation results). We consider two scenarios for the fitness effects of combined compensation: either the combination of both compensatory mutations confers no fitness benefit (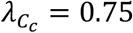, Figure 5A-C) or a high benefit (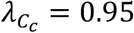, Figure 5D-E) relative to the uncompensated plasmid in the uncompensated chromosomal background 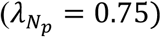. For each simulation we compute three outcomes: plasmid persistence at t_end_, dominant (i.e., largest) population at t_end_, and the succession of dominant plasmid populations (see Methods).

**Figure 5.**
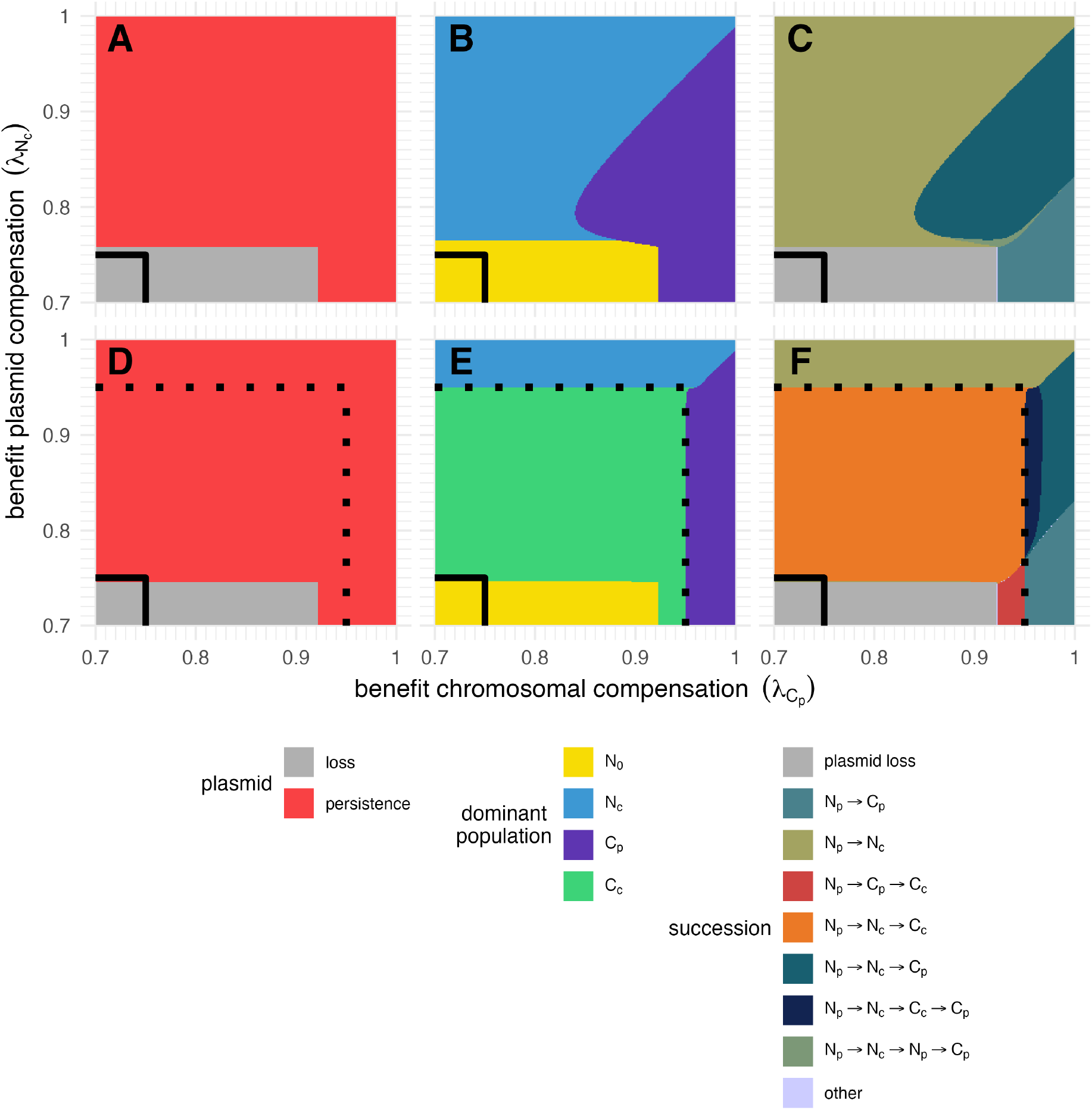
Relative benefits between chromosomal 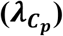, plasmid-borne 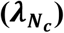, and both compensatory mutations 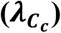 determine whether plasmids persist, which compensatory mutations dominate, and how they succeed each other. Panels **A-C** and **D-F** show different outcomes of the same parameter sweep across 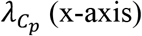 (x-axis) and 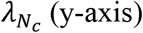 (y-axis), each ranging from 0.7 to 1. Panels **A-C** assume that the combination of both compensatory mutations simultaneously provides no benefit 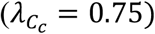 whereas panels **D-F** assume a high benefit 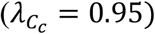 relative to the uncompensated plasmid in the uncompensated chromosomal background benefit 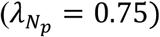. All other parameters are kept constant (Table S1). The solid black line marks the region where both 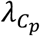 and 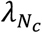 have lower fitness than the uncompensated plasmid 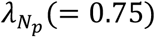. The dotted black line marks where either 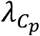 and/or 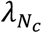 exceed the fitness of 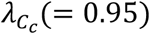. (**A, D**) Persistence is defined as the sum of plasmid-carrying populations exceeding 10^-16^ (numerical zero) at the end of simulation (t_end_ = 10,000h). (**B, E**) The dominant population is the largest population at t_end_. (**C, F**) Succession of transiently dominant plasmid populations can lead to the same final dominant population (shown in **B**). Successions that occur in less than 1% of simulations are grouped as “other” to improve readability. See Methods for a description of the model.

If neither *C*_*p*_ nor *N*_*c*_ have a fitness advantage relative to *N*_*p*_ the plasmid is lost regardless of whether combined compensations confers no or a high benefit (Figure 5A,D). Even though *C*_*c*_ would ensure plasmid persistence, the intermediates *N*_*c*_ and *C*_*p*_ do not accumulate to sufficient sizes for *C*_*c*_ to emerge. Already small benefits of plasmid compensation 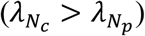 are sufficient for plasmid persistence, but slightly higher benefits are necessary if 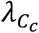 is not beneficial (Figure 5A,D). In contrast, if only *C*_*p*_ has a fitness benefit 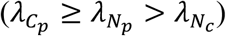 it must be substantially higher to ensure plasmid persistence irrespective of 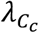(>0.92, Figure 5A,D). This is because only for high benefits can *C*_*p*_ grow fast enough to reach a population size from which *C*_*c*_ can emerge before *t*_*end*_.

If the plasmid persists, the largest population at *t*_*end*_ indicates which location is favoured for compensation (Figure 5B,E). If combined compensation is not beneficial (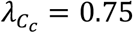, Figure 5A-C), whether *N*_*c*_ or *C*_*p*_ dominates is mainly determined by which benefit is higher. However, *N*_*c*_ can dominate even if its fitness is slightly smaller than that of *C*_*p*_, because horizontal spread allows it to establish faster. As a result, *C*_*p*_ can only overtake *N*_*c*_ by vertical replication before *t*_*end*_ if its growth rate is sufficiently higher. If the plasmid persists, and combined compensation simultaneously confers a benefit (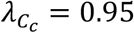, Figure 5D-E), *C*_*c*_ dominates whenever it has the highest growth rate (indicated by black dashed line, Figure 4E) and only if *C*_*p*_ or *N*_*c*_ have the highest growth rate do they dominate instead.

We already illustrated (Figure 4E) that multiple compensatory mutants can spread and intermittently dominate before being succeeded by another. Which succession occurs depends on both the magnitude of compensatory benefits and the location where compensatory mutations emerge (Figure 5C,F). Plasmid compensation *N*_*c*_ typically appears early in successions because of rapid initial horizontal spread, even when it provides only a small benefit, and is ultimately replaced by chromosomal *C*_*p*_ or combined compensation *C*_*c*_ (if these have higher growth rates). In contrast, *C*_*p*_, which cannot spread horizontally, never precedes *N*_*c*_ (Figure 5C,F) and only precedes highly beneficial *C*_*c*_ when plasmid compensation is weak or detrimental (Figure 5F). As a result, when chromosomal compensation is only slightly more beneficial than combined compensation and plasmid compensation is weaker, the succession *N*_*p*_ → *N*_*c*_ → *C*_*c*_ → *C*_*p*_ follows from the horizontal spread advantage of plasmid-borne compensation and the growth rate hierarchy between compensated types (Figure 4C). These findings – that location and the growth rate hierarchy determine succession – are robust and hold across a wide range of parameters, as our sensitivity analyses confirm. Our sensitivity analyses consistently identify growth rates *λ*_*i*_ as the main determinant of which succession occurs, whether growth rates can confer equal benefits across locations (uniform; Figure S10, S12, S14) or are biased according to empirical distributions (gamma; Figure S11, S13, S15), even though these approaches differ in how often plasmids persist and which successions predominate. In summary, initial spread is commonly dominated by plasmid-borne compensation but in the long term plasmid-borne compensation is succeeded by chromosomal or combined compensation when they confer sufficiently higher benefits.

### Trade-offs with resistance and conjugation shape compensatory evolution and successions of plasmid populations differently

Plasmid-encoded functions, such as antibiotic resistance [9,52] or conjugation [27,41,42] can be metabolically expensive. In principle, mutations – arising on the chromosome, plasmid, or both – could reduce these functions and thereby increase growth rates and compensate plasmid fitness costs [11,19,27,53] (Figure S1). To assess the effect of trade-offs, we consider three cases: trade-offs associated with plasmid-borne compensation only (“plasm. only”), chromosomal compensation only (“chrom. only”), or combined (“chrom. & plasm.”). For simplicity, we here consider the scenario of combined compensatory mutations conferring no benefit 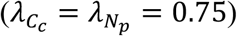 and proportional trade-offs (trade-off strength *p* = 1), i.e., an x% reduction in function corresponds to an x% reduction in cost.

A trade-off between resistance and compensation only becomes relevant in the presence of antibiotics, because in absence of antibiotics resistance provides no benefit. Importantly, antibiotic selection (A=0.9) facilitates persistence of the resistance conferring plasmid independent of compensatory evolution (Figure 6A, no trade-off). Further, since antibiotics remove plasmid-free bacteria, the advantage of horizontal spread for plasmid-borne compensation (Figure 5C,F) is lost, because no recipients for conjugation remain, and whether *N*_*c*_ or *C*_*p*_ dominates is simply determined by which benefit is higher (Figure 6A). However, when compensation is associated with a resistance trade-off, this is no longer the case. Regardless of whether the trade-off affects plasmid and/or chromosomal mutations, for any increase in growth rate the corresponding loss of resistance prevents the affected mutant populations from dominating under antibiotic treatment. Instead, compensation at the unaffected location is favoured, if it confers a sufficient benefit, or the uncompensated ancestral plasmid *N*_*p*_ dominates. Notably, when both chromosome and plasmid are affected by the trade-off, this holds across the full parameter range of benefits. Varying the benefit of combined compensatory mutations and trade-off strengths p does not qualitatively change these results (Figures S17, Figure S18).

**Figure 6.**
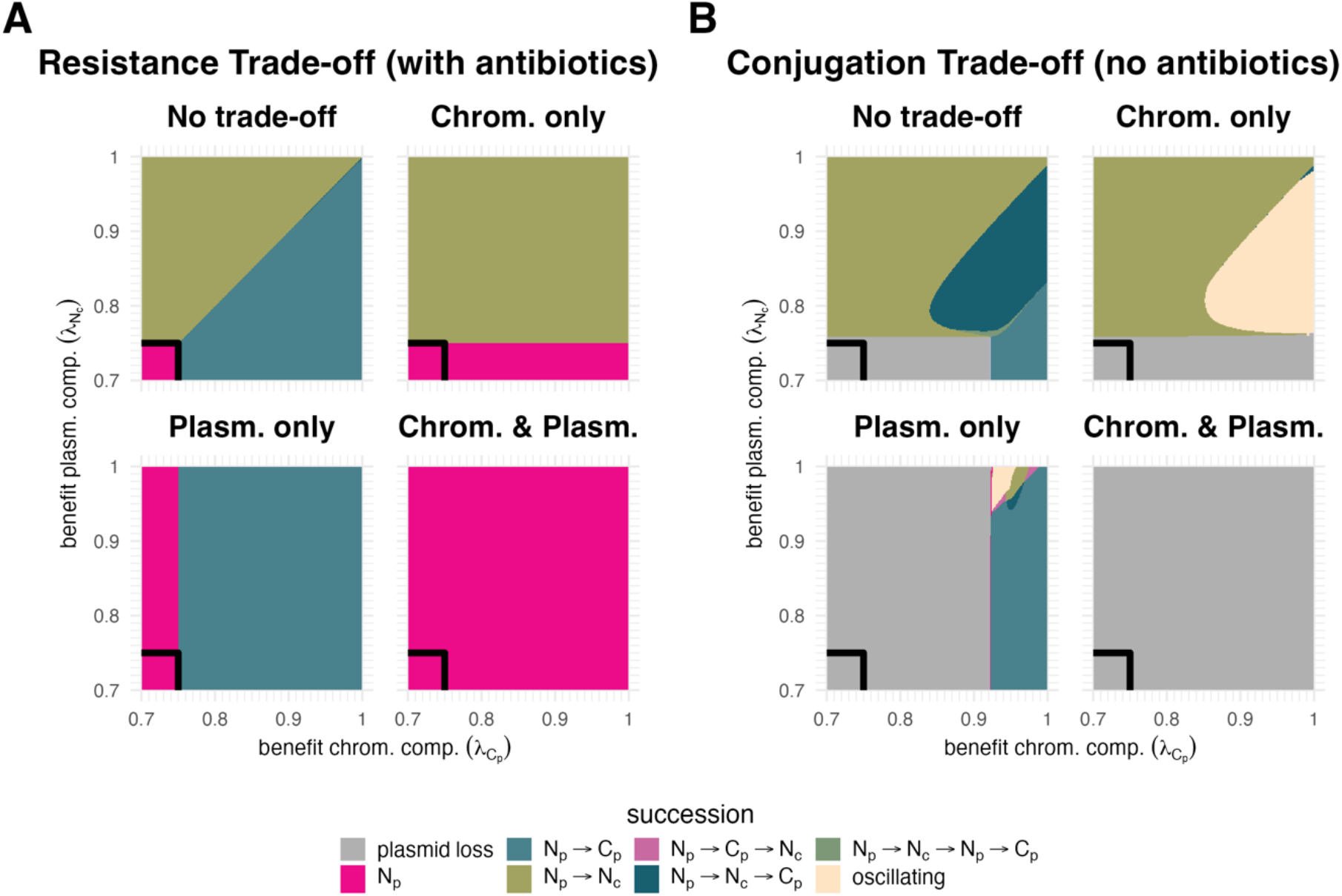
Trade-offs between compensatory benefits and conjugation or resistance constrain or expand the possible successions to the final dominant population. Panels (**A**) and (**B**) show the effects of resistance trade-offs under antibiotic selection and conjugation trade-offs in absence of antibiotics, respectively. Each subpanel shows the succession of transiently dominant plasmid populations across 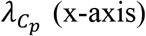 (x-axis) and 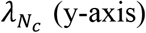 (y-axis), both ranging from 0.7 to 1, corresponding to scenarios without a trade-off (“No trade-off”), with a trade-off associated with mutations located on the chromosome only (“Chrom. only”), on the plasmid only (“Plasm. only”), or on both (“Chrom. and Plasm.”). Conjugation and Resistance trade-off are both assumed to be proportional (i.e., trade-off strength p=1, Methods for details). “Plasmid loss” is defined as <0.1% plasmid-carrying bacteria at end of simulation (t_end_ = 10,000h). “Oscillating” describes successions where the same populations replace each other repeatedly. Lastly, “other” summarises successions occurring in less than 1% of simulations.

Next, we investigate the effects of a trade-off between growth rate compensation and conjugation rate (in the absence of antibiotics). Without trade-offs, we observe sequential successions (e.g. *N*_*p*_ → *N*_*c*_ → *C*_*p*_ or *N*_*p*_ → *N*_*c*_, Figure 5C). Introducing a conjugation trade-off changes these dynamics qualitatively. Because compensatory benefits and conjugation reduction are proportional (trade-off strength p = 1), even small benefits can reduce conjugation below the rate necessary for plasmid maintenance. If the conjugation trade-off is only associated with chromosomal mutations, chromosomal compensation alone cannot prevent plasmid loss, but the plasmid can still persist through plasmid compensation (if 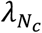 is high enough, Figure 6B “Chrom. Only”). If instead only plasmid mutations are associated with the trade-off, the plasmid is lost across a broader range of benefits (Figure 6B “Plasm. Only”). Here, persistence relies solely on chromosomal compensation, which generally requires higher benefits 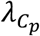 because it cannot spread horizontally. If mutations in either location come with the trade-off, no unpenalized compensation remains and the plasmid is lost for all combinations of benefits.

With a conjugation trade-off, we also observe interesting new dynamics: ‘oscillations’, where the same populations replace each other repeatedly (Figure 6B, see Figure S19 for examples). This happens mainly in the chromosome-only and, more rarely, the plasmid-only scenario. The oscillations stem from frequency-dependent selection caused by differences in conjugation rate. For example, in Figure 6B “Chrom. only”, *N*_*c*_ initially spreads via conjugation but as *N*_0_ is depleted this fitness advantage wanes, allowing *C*_*p*_, which has a higher growth rate (below the diagonal), to increase in frequency. However, due to the chromosomal conjugation trade-off *C*_*p*_ cannot maintain its plasmid, giving rise to *C*_0_. Since *C*_0_ bears the cost of chromosomal compensation in the absence of plasmid infection, it is outcompeted by *N*_0_, restoring the conditions for *N*_*c*_ to spread and the cycle repeats. Therefore, neither chromosomal nor plasmid-borne compensation gains a lasting advantage, and no single compensatory location is clearly favoured for these benefit combinations. These patterns hold qualitatively for different trade-off strengths p and when combined compensation 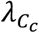 is highly beneficial (Figure S20, Figure S21). Overall, we find that trade-offs between plasmid cost compensation and resistance or conjugation traits can constrain the spread of compensatory mutations in all locations, but they can also lead to oscillating succession of compensated and un-compensated populations.

## Discussion

In this study we use mathematical modelling to answer whether mutations that compensate for plasmid carriage costs are expected to be located on the plasmid, the chromosome, or both. Our main finding is that the ultimate, long-term location of compensation is driven by the fitness benefit conferred, and not by the location per se. Using analytical invasion analysis (Figure 3, Table S5) and numerical simulations (Figure 4-6) we show that whether a given type of compensated mutation can be invaded by another type depends primarily on the growth rates (*λ*_*i*_) as long as compensation does not trade off with either conjugation or resistance. However, the locations of the compensatory mutations shape succession dynamics, as plasmid borne compensation tends to rise faster initially even when eventually replaced by chromosomal compensation (Figure 5, Figure S10-12).

To characterise how costs and benefits of mutations vary across locations, we reviewed existing data from plasmid evolution experiments (Figure 2, see Methods for details). These data suggest that combined compensation confers the highest average benefit. However, these data are likely confounded by chromosomal mutations that are beneficial regardless of plasmid carriage, e.g. through adaptation to the experimental setup [54,55]. Distinguishing strictly compensatory chromosomal mutations [49] from generally beneficial ones requires comparing fitness across all combinations of ancestral/evolved chromosomes and plasmids, which only some studies have done [13,19,29,45,48]. Where the data allowed, we excluded non-compensatory beneficial mutations, yet combined compensation still showed the highest average benefit (Figure 2, lower violin plots; too few samples remain when restricted to strictly compensatory mutations). Given this bias and methodological differences between studies, we use these data mainly to inform the overall range of the population growth rates *λ*_*i*_ in our model parameterisation (Figure 5,6, Table S6). Nonetheless, these data suggest that compensatory benefits differ among locations within experiments but span similar ranges across experiments.

Arguments for either plasmid-borne or chromosomal compensatory mutations being ‘superior’ have been made both on theoretical and experimental grounds. The main argument for plasmid-borne compensation is horizontal spread: Zwanzig et al. (2019) showed via mathematical modelling that, when each location is considered in isolation, plasmid-borne compensation facilitates plasmid survival while chromosomal compensation with the same benefit cannot. Accordingly, we find that when each location is considered in isolation (*μ*_*c*_ = 0 or *μ*_*p*_ = 0), plasmid-borne compensation establishes faster and enables plasmid survival more often than chromosomal compensation (Figure 4, Figure S9). However, when both types of compensation can emerge (*μ*_*p*_ ≠ 0 and *μ*_*c*_ ≠ 0), plasmid-borne compensation rises fast initially, but can eventually be replaced by chromosomal or combined compensation (Figure 5, Figure S8).

The argument for chromosomal compensation being superior is that plasmid transfer from chromosomally compensated cells to uncompensated competitors reduces the recipient’s fitness sufficiently such that donors can outcompete them – a process coined plasmid “weaponization” [39,39,40,51]. We find that “weaponization” is mainly relevant under low segregation loss and low cost of chromosomal compensation in absence of plasmid (Figures S3-5). Generally, “weaponization” is most effective when plasmid spread is self-sustained in compensated donor and uncompensated competitor populations (basic reproductive number, *R*_0_ > 1 [56]) – which both conditions contribute to – as otherwise too few competitors receive the plasmid. Interestingly, we find that chromosomal compensation can benefit from the prior spread of plasmid-borne compensation. While the uncompensated plasmid cannot persist (*R*_0_ < 1), the compensated plasmid can (*R*_0_ > 1) and its spread reduces competing plasmid-free cells, allowing chromosomal compensation – which confers a higher growth rate – to establish faster than if the compensated but still costly plasmid had not already spread, effectively co-opting plasmid “weaponization” without contributing itself (Figures S7-9).

Notably, self-sustained spread likely matters less in spatially structured environments [39,51], where spread to close neighbours already provides a local competitive benefit without requiring secondary infections.

Our model includes combined compensation, the simultaneous presence of chromosomal and plasmid-borne compensation in the same cell, which prior studies have not focused on. Unlike plasmid-borne compensation, mutants with combined compensation cannot increase in population size by conjugation, as this would require a chromosomally compensated plasmid-free recipient (*C*_0_, Figure 1), which is virtually absent (Figure 3). Like chromosomal compensation, it can benefit from plasmid “weaponization”, but only under the same conditions (low segregation loss and costs of chromosomal compensation, see above). Uniquely, combined compensation depends on either single mutant reaching sufficient frequency first and therefore emerges slowly when it provides little additional benefit (Figure 3) or not at all when both single mutants are detrimental (Figure 5). Yet combined compensation could still be “superior” if it confers the highest benefit (Figure 3, Figure 5D-F) – which empirical data suggest, and which is also expected if the benefits of the single compensations are additive.

When plasmid costs are linked to the expression of ARGs or the conjugation machinery, compensatory mutations that ameliorate these costs by reducing expression can generate trade-offs. We explored how the location and strength of such trade-offs can shape compensatory evolution (see Methods). For trade-offs between compensation and resistance, our simulations show that the location of compensation plays little role, as plasmid-free recipients are susceptible and removed by the antibiotic, eliminating horizontal spread as the main source of difference between locations (Figure 6A, Figure S17-18). Conjugation trade-offs, in contrast, do show location dependence: plasmid-associated trade-offs cause more plasmid loss, whereas chromosome-associated trade-offs have less impact, as establishment of chromosomal compensation benefits less from horizonal spread (Figure 6B, Figure S20-21). For both resistance and conjugation trade-offs, sufficiently strong trade-offs prevent compensation, as reduced resistance increases antibiotic-induced death more than the compensatory gain in growth, or cause plasmid loss when conjugation is reduced too much. As resistance and conjugation machinery are plasmid-encoded, compensating their costs on the plasmid may more often be associated with trade-offs [11,40,47,57], which could shift compensation toward the chromosome and explain why chromosomal compensation is more frequently observed [19,25,40].

Our model makes several simplifying assumptions that may limit generalisability. First, our simulations are deterministic, implying that populations cannot go extinct in finite time but persist at arbitrarily small sizes, and can still give rise to new mutants. This corresponds to a biological scenario where most genotypes are continuously present by recurrent immigration or (back-) mutations in large populations. In finite populations, however, rare genotypes can stochastically go extinct before acquiring further mutations. This could make plasmid-borne compensation harder to replace once dominant, as the genotypes that could replace it may no longer emerge once their precursors are depleted. Second, we assume effectively one plasmid copy per cell and no co-infection by compensated and uncompensated plasmids. This is reasonable for conjugative plasmids which typically have low copy numbers [58,59], and carry exclusion systems that prevent co-infection [see 60, and 61 for review]. However, higher copy numbers can accelerate or hinder the emergence of plasmid-borne compensation by increasing effective mutation rates or via segregational drift, respectively [62–64]. Finally, we assume a constant environment. In natural and clinical settings, nutrient availability fluctuates and can shape selection – which can affect plasmid costs or conjugation rates [65–68]. In the clinic, fluctuating antibiotic exposure may allow compensation with resistance trade-offs during drug-free periods but select against it under treatment. How fluctuating environments shape which compensation type establishes remains an open question for future work.

What do our findings mean for the spread and persistence of plasmids and AMR? In the short term, under little or no treatment and without trade-offs – conditions typical for plasmid spread in commensals – plasmid-borne compensation can pose a more urgent public-health threat, since it drives fast short-term spread of AMR plasmids. In the longer term, however, chromosomal or combined compensation may take over, potentially giving rise to bacterial hosts with the capacity to sustain multiple AMR plasmids.

## Acknowledgments

We thank S. Lehtinen, B. Siedentop and C. Igler for helpful discussions and valuable feedback on the manuscript. We used Large Language Models (Claude Sonnet 4.6 and Opus 4.8; ChatGPT 4o, o1, and 5.5) for assistance with coding (including Claude Code) and language editing of this manuscript.

## Notes

### Competing Interest Statement

The authors have declared no competing interest.

