## Supplemental Information for "Chromosome, Plasmid, or Both: Short-Term Dynamics and Long-Term Outcomes of Plasmid Cost Compensation under Trade-Offs"

Text S1. **Differential Equations.**

$$
\begin{aligned}
\frac{\mathrm{d}N_0(t)}{\mathrm{d}t} = {} & (1-\mu_c)\lambda_{N_0}N_0(t) + s\Big((1-\mu_c)\lambda_{N_c}N_c(t) + \big(1-\mu_p-\mu_c\big)\lambda_{N_p}N_p(t)\Big) \\
& - \beta\left(N_p(t) + r_pN_c(t) + r_cC_p(t) + r_cr_pC_c(t)\right)N_0 - AN_0(t) \\
& - \theta(t)N_0(t) \qquad [\boldsymbol{Eqn.}\,\mathbf{1}]
\end{aligned}
$$

$$
\begin{aligned}
\frac{\mathrm{d}N_p(t)}{\mathrm{d}t} = {} & \big(1-\mu_p-\mu_c\big)\lambda_{N_p}N_p(t) - s\big(1-\mu_p-\mu_c\big)\lambda_{N_p}N_p(t) \\
& + \beta\Big(\big(1-\mu_p\big)N_p(t)N_0(t) + \big(1-\mu_p\big)r_cC_p(t)N_0(t)\Big) - (1-\rho)AN_p(t) \\
& - \theta(t)N_p(t) \qquad [\boldsymbol{Eqn.}\,\mathbf{2}]
\end{aligned}
$$

$$
\begin{aligned}
\frac{\mathrm{d}N_c(t)}{\mathrm{d}t} = {} & (1-\mu_c)\lambda_{N_c}N_c(t) + \mu_p\lambda_{N_p}N_p(t) + \beta\left(r_pN_c(t)N_0(t) + r_cr_pN_0(t)C_c(t)\right) \\
& + \mu_p\beta\left(N_p(t) + r_cC_p(t)\right)N_0 - \big(1-\rho(1-l_p)\big)AN_c(t) - s(1-\mu_c)\lambda_{N_c}N_c(t) \\
& - \theta(t)N_c(t) \qquad [\mathbf{Eqn.\,3}]
\end{aligned}
$$

$$
\begin{aligned}
\frac{\mathrm{d}C_0(t)}{\mathrm{d}t} = {} & \lambda_{C_0}C_0(t) + \mu_c\lambda_{N_0}N_0(t) + s\left(\lambda_{C_c}C_c(t) + \big(1-\mu_p\big)\lambda_{C_p}C_p(t)\right) \\
& - \beta\left(N_p(t) + r_pN_c(t) + r_cC_p(t) + r_cr_pC_c(t)\right)C_0(t) - AC_0(t) \\
& - \theta(t)C_0(t) \qquad [\boldsymbol{Eqn.}\,\mathbf{4}]
\end{aligned}
$$

$$
\begin{aligned}
\frac{\mathrm{d}C_p(t)}{\mathrm{d}t} = {} & \lambda_{C_p}C_p(t) + \mu_c\lambda_{N_p}N_p(t) - s\big(1-\mu_p\big)\lambda_{C_p}C_p(t) \\
& + \beta\big((1-\mu_p)N_p(t)C_0(t) + r_c(1-\mu_p)C_p(t)C_0(t)\big) - (1-\rho(1-l_c))AC_p(t) \\
& - \theta(t)C_p(t) \qquad [\boldsymbol{Eqn.}\,\mathbf{5}]
\end{aligned}
$$

$$
\begin{aligned}
\frac{\mathrm{d}C_c(t)}{\mathrm{d}t} = {} & \lambda_{C_c}C_c(t) + \mu_p\lambda_{C_p}C_p(t) + \mu_c\lambda_{N_c}N_c(t) + \mu_p\beta\left(N_p(t) + r_cC_p(t)\right)C_0(t) \\
& + \beta\left(r_pN_c(t) + r_cr_pC_c(t)\right)C_0(t) - s\lambda_{C_c}C_c(t) \\
& - \left(1-\rho\left(1-\big(l_c+l_p-l_cl_p\big)\right)\right)AC_c(t) \\
& - \theta(t)C_c(t) \qquad [\boldsymbol{Eqn.}\,\mathbf{6}]
\end{aligned}
$$

$\theta(t)$ is defined such that $\sum\frac{dX_i}{dt} = 0$, with $X \epsilon \{N, C\}$ and $i \epsilon \{0, p, c\}$.

Text S2 **Sensitivity and robustness analyses.**

To test whether our findings – mainly that plasmid-borne compensation $N_c$ emerges first and that $C_p$ and $C_c$ are initially disadvantaged but can become dominant later – hold across a wider range of parameters we sampled 100,000 parameter sets using two complementary approaches: "gamma" and "conditional uniform". The "gamma" approach samples growth rates $\lambda_i$ from gamma distributions fitted directly to the empirical data (excluding cases where $\lambda_{C_0} > 1$, see Results for rationale). For "conditional uniform", growth rates $\lambda_i$ are sampled uniformly from ranges informed by the empirical fitness effects but ignoring tails (Figure 2), retaining only parameter sets that contain at least one compensation type that is beneficial, – i.e. either $N_c, C_p$, or $C_c$ has a higher growth rate than $N_p$ – until 100,000 sets are reached. For both sampling schemes, the segregation loss rate $s$ and the conjugation rate $\beta$ are sampled log-uniformly from biologically realistic ranges based on the literature [1–3]. All other parameters are not varied unless explicitly stated. See Table S6 for an overview of values used. Together, these two approaches allow us to test whether our findings are robust to when the benefits of compensatory mutations are biased per location and when, a priori, there is no difference in possible fitness effects between locations.

We then run simulations for each sampled parameter set and categorize the outcome by plasmid persistence, dominant population and succession (see Methods for descriptions). To identify which parameters distinguish the different succession outcomes, and thereby where compensatory mutations should establish, we use two supervised classification methods: Linear Discriminant Analysis (LDA) and eXtreme Gradient Boosting (XGBoost). LDA is a method based on dimension reduction. Intuitively, an LDA takes a multi-dimensional dataset – in our case the dimensions correspond to the number of varied parameters – and projects it to a lower dimensional subspace so that the categorical outcome or classes are maximally separated. The contribution of each parameter to this class separation is given by its discriminant coefficient and can be depicted as arrows, whose length and angle indicate the magnitude and direction of its contribution to the separation. LDA provides an easily interpretable picture of parameter importance and directionality for the outcome and often performs reasonably well even for non-normally distributed data (as in our uniform sampling regime) [4]. However, LDA cannot identify non-linear effects well if present. To address these limitations, we also use XGBoost. XGBoost is based on gradient-boosted trees, in which an ensemble of decision trees is trained sequentially, with each new tree correcting the errors of the current model, thereby boosting the classification performance of the ensemble. To interpret the XGBoost model predictions we compute Shapley values. Briefly, Shapley values quantify each parameter´s contribution to the model predictions (here the tree ensemble), weighted and summed over all possible combinations of parameter values. Jointly, XGBoost and Shapley values make it possible to interpret how important each parameter is for the outcome and whether increasing that parameter increases or decreases its likelihood. LDA and XGBoost and Shapley value analyses were conducted in *R* (v. 4.5.0) using the packages *MASS*, *xgboost*, and *shapviz*. For convenience, we utilized *RCall.jl* (v. 0.14.8) for passing simulation results between *Julia* and *R*.

To assess specifically the temporal dynamics of beneficial compensation across a wide range of parameters, we additionally vary a single focal growth rate systematically over 21 values spanning empirically realistic ranges (≈0.76–0.99), while keeping $\lambda_{N_p}$ fixed to 0.75. All other parameters are uniformly sampled by Latin Hypercube sampling (10,000 sets per focal value, see Table S6 for values) to ensure efficient and comprehensive coverage of the parameter space [4]. Briefly, in LHS, each parameter range is divided into a set number of intervals, and each interval is sampled uniformly once. We consider three scenarios: $N_c$ or $C_p$ in isolation ($\mu_c$ or $\mu_p = 0$); $N_c$ and $C_p$ in competition, with $C_c$ "non-beneficial" ($\lambda_{C_c}$=$\lambda_{N_p}$); or $N_c$, $C_p$ and $C_c$, competing against each other. We vary $\lambda_{N_c}$ and $\lambda_{C_p}$ as focal parameters across all three scenarios. $\lambda_{C_c}$ is only varied as a focal parameter in the scenario with competition among all three types – as it is set to be "non-beneficial" in the others. For each focal growth rate value, among the simulations in which the plasmid persisted, we record (i) the proportion of runs in which the focal compensated population is the dominant surviving plasmid population at the end and (ii), among those runs, the median time to dominance (time to reach 50% of the total population). This allows us to compare both how often and how quickly plasmid versus chromosomal compensation prevails as a function of its fitness benefit.

Text S3 **Detailed Description of the analytical invasion analysis.**

*Without segregation loss, mutations, and trade-offs, all locations are possible, but chromosome and dual compensation are overrepresented*

To assess what determines where compensatory mutations are expected to be located – on the plasmid, the chromosome, or on both – we first analytically investigate a simplified system, where we assume that segregation loss and mutation rates are negligible ($\mu_c, \mu_p, l_c, l_p, s = 0,$ and no trade-offs $l_c, l_p, s = 0,\ r_c, r_p = 1$). We find that for this simplified system a total of ten equilibria are biologically feasible: six correspond to each population existing alone ($N_0, N_p, N_c, C_0, C_p, C_c$) and four represent stable coexistence of three populations ($N_0, N_c, C_c$; $N_0, N_p, C_p$; $N_c, C_0, C_c$; and $N_p, C_0, C_p$) (Figure 3, Table S5 for details). In two co-existence states, compensation is located only on the chromosome ($N_0, N_p, C_p$ and $N_p, C_0, C_p$), while in the other two ($N_0, N_c, C_c$ and $N_c, C_0, C_c$) populations with compensatory mutations on both the chromosome and plasmid ($C_c$) and only on the plasmid ($N_c$) co-exist (Figure 3B,C). However, equilibria where both the compensated ($N_p$ or $C_p$) and uncompensated ($N_c$ or $C_c$) plasmids stably coexist are not possible for this simplified system. Because both plasmids have identical conjugation and segregation loss rates, the lower-cost plasmid outcompetes the other. Notably, two population equilibria are not possible under turbidostat death but can occur under alternative mortality assumptions. Three population equilibria are not biologically feasible for all parameter conditions (see Table S5), where feasibility implies that all six populations are larger or equal to zero.

*Compensatory mutations are stable where benefits are highest*

We further determine under which parameter conditions equilibria can successfully be invaded by any of the other populations (Figure 3 for examples and Table S4 for details). Equilibria of plasmid-free populations ($N_0$ or $C_0$) can be invaded by plasmid carriers ($N_p, N_c, C_p$, or $C_c$) if the sum of antibiotic pressure ($A$), conjugation rate ($\beta$) and the growth rate of the invading population ($\lambda_i$) is larger than the growth rate of the plasmid-free population, i.e., $A + \beta + \lambda_i > \lambda_{N_0,C_0}$, where $\lambda_i \in \{\lambda_{N_p}, \lambda_{N_c}, \lambda_{C_p}, \lambda_{C_c}\}$. Conversely, plasmid-free bacteria can invade plasmid-carriers if their growth rate is sufficiently high, i.e., $\lambda_{N_0,C_0} > A + \beta + \lambda_i$ (Figure 3A). Moreover, plasmid-free bacteria, i.e., $N_0$ or $C_0$, can invade the other only if they have a higher growth rate ($\lambda_i$). Similarly, plasmid carrying populations ($N_p, N_c, C_p$, or $C_c$), can be invaded by another plasmid carrying population only if the invader has a higher growth rate ($\lambda_i$) (Figure 3A). For the three population equilibria, invasion by any of the absent populations is determined by competition between the invader and a single resident population. Specifically, invasion occurs if the population of singular chromosome type present at the equilibrium has a lower growth rate than the invader of the same chromosome type (Figure 3B). Additionally, each equilibrium can be invaded if the present plasmid free population, $N_0$ or $C_0$, has a lower fitness than the same chromosome type with the absent plasmid type, i.e., $A+\ \beta+\lambda_i>\lambda_{N_0,C_0}$. Lastly, the absent plasmid-free populations, $N_0$ or $C_0$, can invade if they have a higher growth rate than the present plasmid-free population – i.e., if chromosomal compensation without plasmid carriage is costly or beneficial (Figure 3B).

Table S1. **Model parameters and default values for numerical simulations.**

| **Parameter** | **Description** | **Value** | **Unit** |
|---|---|---|---|
| $\lambda_{N_0}$ | Growth rate $N_0$ | 1 | $h^{-1}$ |
| $\lambda_{N_p}$ | Growth rate $N_p$ | 0.75 | $h^{-1}$ |
| $\lambda_{N_c}$ | Growth rate $N_c$ | 0.95 | $h^{-1}$ |
| $\lambda_{C_0}$ | Growth rate $C_0$ | 0.999 | $h^{-1}$ |
| $\lambda_{C_p}$ | Growth rate $C_p$ | 0.95 | $h^{-1}$ |
| $\lambda_{C_c}$ | Growth rate $C_c$ | 0.95 | $h^{-1}$ |
| $s$ | Segregation loss rate | 0.01 | $h^{-1}$ |
| $\beta$ | Conjugation rate | 0.25 | $\frac{CFU}{mL\ h^{-1}}$ |
| $\mu_c$ | Mutation rate of chromosome | $1.5\times10^{-5}$ | / |
| $\mu_p$ | Mutation rate of plasmid | $1.5\times10^{-5}$ | / |
| $A$ | Killing rate of antibiotic | 0 (1 in Figure 6A, S17, S18) | $h^{-1}$ |
| $\rho$ | Reduction of antibiotic killing by plasmid-conferred resistance | 0.9 | / |
| $l_c$ | Reduction of resistance by chromosomal compensation | 0 | / |
| $l_p$ | Reduction of resistance by plasmid compensation | 0 | / |
| $r_c$ | Fraction of conjugation remaining under chromosomal compensation | 1 | / |
| $r_p$ | Fraction of conjugation remaining under plasmid compensation | 1 | / |

Table S2**. Summary statistics of plasmid costs and benefits of compensatory mutations extracted from the literature.** Mean fitness values (bootstrapped 95% confidence intervals) and minimum and maximum reported relative fitness measures, for the full dataset and when sets of fitness measures where $\lambda_{C_0} > 1$ are excluded (see Results for rationale). All fitness measures are normalized to the plasmid free ancestor ($\lambda_{N_0} = 1$).

| **Locations of compensation** | **Means All Data (95%CI)** | **Range All Data (min, max)** | **Means $\lambda_{C_0} > 1$ excluded (95%CI)** | **Range $\lambda_{C_0} > 1$ excluded (min, max)** |
|---|---|---|---|---|
| $\lambda_{C_c}$ | 1.128 (1.044, 1.211) | (0.880, 2.150) | 1.013 (0.977, 1.048) | (0.880, 1.192) |
| $\lambda_{C_p}$ | 1.012 (0.943, 1.081) | (0.526, 1.302) | 0.926 (0.839, 1.014) | (0.526, 1.100) |
| $\lambda_{C_0}$ | 1.367 (1.153, 1.581) | (0.940, 2.370) | 0.970 (0.954, 0.987) | (0.940, 0.990) |
| $\lambda_{N_c}$ | 0.974 (0.906, 1.042) | (0.697, 2.051) | 0.958 (0.880, 1.035) | (0.697, 2.051) |
| $\lambda_{N_p}$ | 0.776 (0.719, 0.832) | (0.360, 0.970) | 0.743 (0.670, 0.816) | (0.360, 0.943) |

Table S3. **Plasmid cost and benefits for only head-to-head/direct competition studies.** Mean fitness values (bootstrapped 95% confidence intervals) and minimum and maximum reported relative fitness measures, if sets with fitness measures where $\lambda_{C_0} > 1$ are excluded or not (see Results for rationale). Studies reporting Malthusian ratios from competition assays are [5–15].

| **Locations of compensation** | **Means All Data (95%CI)** | **Range All Data (min, max)** | **Means $\lambda_{C_0} > 1$ excluded (95%CI)** | **Range $\lambda_{C_0} > 1$ excluded (min, max)** |
|---|---|---|---|---|
| $\lambda_{C_c}$ | 1.136 (1.047, 1.225) | (0.880, 2.150) | 1.013 (0.973, 1.052) | (0.880, 1.192) |
| $\lambda_{C_p}$ | 1.033 (0.959, 1.108) | (0.526, 1.302) | 0.949 (0.834, 1.063) | (0.526, 1.100) |
| $\lambda_{C_0}$ | 1.423 (1.187, 1.658) | (0.940, 2.370) | 0.966 (0.942, 0.989) | (0.940, 0.990) |
| $\lambda_{N_c}$ | 0.974 (0.906, 1.042) | (0.697, 2.051) | 0.958 (0.880, 1.035) | (0.697-2.051) |
| $\lambda_{N_p}$ | 0.766 (0.705, 0.827) | (0.360, 0.970) | 0.722 (0.642, 0.803) | (0.360, 0.943) |

Table S4. **Summary statistics of Plasmid costs and benefits of compensatory mutations based on study-level mean fitness measures.** In contrast to Table S3, fitness measures were first summarized as one mean value per study across all reported evolved lines or strains. Means across these study-level values and bootstrapped 95% confidence intervals were then calculated. Means, minimum and maximum values are reported both with and without excluding datasets in which $\lambda_{C_0} > 1$ (see Results for rationale).

| **Locations of compensation** | **Means All Data (95%CI)** | **Range All Data (min, max)** | **Means $\lambda_{C_0} > 1$ excluded (95%CI)** | **Range $\lambda_{C_0} > 1$ excluded (min, max)** |
|---|---|---|---|---|
| $\lambda_{C_c}$ | 1.123 (1.004, 1.242) | (0.931, 1.583) | 1.058 (0.976, 1.140) | (0.932, 1.192) |
| $\lambda_{C_p}$ | 0.958 (0.811, 1.105) | (0.526, 1.260) | 0.901 (0.738, 1.065)) | (0.526, 1.092) |
| $\lambda_{C_0}$ | 1.220 (0.875, 1.564) | (0.967, 2.250) | 0.973 (0.954, 0.993) | (0.945, 0.990) |
| $\lambda_{N_c}$ | 0.920 (0.8253, 1.020) | (0.748, 1.234) | 0.921 (0.818, 1.023) | (0.748, 1.180) |
| $\lambda_{N_p}$ | 0.769 (0.687, 0.850) | (0.522, 0.917) | 0.707 (0.575, 0.840) | (0.360, 0.910) |

**Table S5. Analytical Invasion Analysis.** Biologically feasible equilibria of a simplified model and parameter conditions for successful invasion by other populations.

| Invaded by \ Equilibrium | $\boldsymbol{N_0}$ | $\boldsymbol{N_p}$ | $\boldsymbol{N_c}$ | $\boldsymbol{C_0}$ | $\boldsymbol{C_p}$ | $\boldsymbol{C_c}$ |
|---|---|---|---|---|---|---|
| $\boldsymbol{N_0}$ | / | $\lambda_{N_0} > A + \beta + \lambda_{N_p}$ | $\lambda_{N_0} > A + \beta + \lambda_{N_c}$ | $\lambda_{N_0} > \lambda_{C_0}$ | $\lambda_{N_0} > A + \beta + \lambda_{C_p}$ | $\lambda_{N_0} > A + \beta + \lambda_{C_c}$ |
| $\boldsymbol{N_p}$ | $A + \beta + \lambda_{N_p} > \lambda_{N_0}$ | / | $\lambda_{N_p} > \lambda_{N_c}$ | $A + \lambda_{N_p} > \lambda_{C_0}$ | $\lambda_{N_p} > \lambda_{C_p}$ | $\lambda_{N_p} > \lambda_{C_c}$ |
| $\boldsymbol{N_c}$ | $A + \beta + \lambda_{N_c} > \lambda_{N_0}$ | $\lambda_{N_c} > \lambda_{N_p}$ | / | $A + \lambda_{N_c} > \lambda_{C_0}$ | $\lambda_{N_c} > \lambda_{C_p}$ | $\lambda_{N_c} > \lambda_{C_c}$ |
| $\boldsymbol{C_0}$ | $\lambda_{C_0} > \lambda_{N_0}$ | $\lambda_{C_0} > A + \beta + \lambda_{N_p}$ | $\lambda_{N_0} > A + \beta + \lambda_{N_c}$ | / | $\lambda_{C_0} > A + \beta + \lambda_{C_p}$ | $\lambda_{C_0} > A + \beta + \lambda_{C_c}$ |
| $\boldsymbol{C_p}$ | $A + \lambda_{C_p} > \lambda_{N_0}$ | $\lambda_{C_p} > \lambda_{N_p}$ | $\lambda_{C_P} > \lambda_{N_c}$ | $A + \beta + \lambda_{C_p} > \lambda_{C_0}$ | / | $\lambda_{C_p} > \lambda_{C_c}$ |
| $\boldsymbol{C_c}$ | $A + \lambda_{C_c} > \lambda_{N_0}$ | $\lambda_{C_c} > \lambda_{N_p}$ | $\lambda_{C_c} > \lambda_{N_c}$ | $A + \beta + \lambda_{C_c} > \lambda_{C_0}$ | $\lambda_{C_c} > \lambda_{C_p}$ | / |

| Invaded by \ Equilibrium | $\boldsymbol{N_0\ N_c\ C_c}$ | $\boldsymbol{N_0\ N_p\ C_p}$ | $\boldsymbol{N_c\ C_0\ C_c}$ | $\boldsymbol{N_p\ C_0\ C_p}$ |
|---|---|---|---|---|
| $\boldsymbol{N_0}$ | / | / | $\lambda_{N_0} > \lambda_{C_0}$ | $\lambda_{N_0} > \lambda_{C_0}$ |
| $\boldsymbol{N_p}$ | $A + \beta + \lambda_{N_p} > \lambda_{N_0}$ | / | $\lambda_{N_p} > \lambda_{N_c}$ | / |
| $\boldsymbol{N_c}$ | / | $A + \beta + \lambda_{N_c} > \lambda_{N_0}$ | / | $\lambda_{N_c} > \lambda_{N_p}$ |
| $\boldsymbol{C_0}$ | $\lambda_{C_0} > \lambda_{N_0}$ | $\lambda_{C_0} > \lambda_{N_0}$ | / | / |
| $\boldsymbol{C_p}$ | $\lambda_{C_p} > \lambda_{C_c}$ | / | $A + \beta + \lambda_{C_p} > \lambda_{C_0}$ | / |
| $\boldsymbol{C_c}$ | / | $\lambda_{C_c} > \lambda_{C_p}$ | / | $A + \beta + \lambda_{C_c} > \lambda_{C_0}$ |

Table S6. **Parameter ranges used for sensitivity and robustness analyses.** Shown are the parameters, their descriptions, and the sampling ranges for conditional uniform and gamma (see Text S2). Segregation loss (s) and conjugation rate ($\beta$) – marked by * – are sampled uniformly on a $\log_{10}$ scale and then back transformed via $10^x$, with ranges shown as actual (back-transformed) values. For sampling, normal distributions are truncated at zero to ensure biologically meaningful values (non-negative) of growth rates ($\lambda_i$). Trade-off presence, type (location, and resistance or conjugation), and strength (p) are set separately.

| **Parameter** | **Description** | **Conditional Uniform (min, max)** | **Gamma (shape, rate)** |
|---|---|---|---|
| $\lambda_{N_0}$ | Growth rate $N_0$ | 1 (fixed) | 1 (fixed) |
| $\lambda_{N_p}$ | Growth rate $N_p$ | 0.74, 1 | 15.39, 20.27 |
| $\lambda_{N_c}$ | Growth rate $N_c$ | 0.74, 1 | 25.88, 27.03 |
| $\lambda_{C_0}$ | Growth rate $C_0$ | 0.74, 1 | 1856.04, 1912.88 |
| $\lambda_{C_p}$ | Growth rate $C_p$ | 0.74, 1 | 31.79, 34.32 |
| $\lambda_{C_c}$ | Growth rate $C_c$ | 0.74, 1 | 108.57, 107.23 |
| $s*$ | Segregation loss rate | $10^{-6}$, 0.1 | $10^{-6}$, 0.1 |
| $\beta*$ | Conjugation rate | $10^{-4}$, 0.25 | $10^{-4}$, 0.25 |
| $\mu_c$ | Mutation rate of chromosome | $1.5\times10^{-5}$ (fixed) | $1.5\times10^{-5}$ (fixed) |
| $\mu_p$ | Mutation rate of plasmid | $1.5\times10^{-5}$ (fixed) | $1.5\times10^{-5}$ (fixed) |
| $A$ | Killing rate of antibiotic | 0 (fixed) | 0 (fixed) |
| $\rho$ | Reduction of antibiotic killing by plasmid-conferred resistance | 0.9 (fixed) | 0.9 (fixed) |
| $l_c$ | Reduction of resistance by chromosomal compensation | 0 (fixed) | 0 (fixed) |

| | | | |
|---|---|---|---|
| $l_p$ | Reduction of resistance by plasmid compensation | 0 (fixed) | 0 (fixed) |
| $r_c$ | Fraction of conjugation remaining under chromosomal compensation | 1 (fixed) | 1 (fixed) |
| $r_p$ | Fraction of conjugation remaining under plasmid compensation | 1 (fixed) | 1 (fixed) |

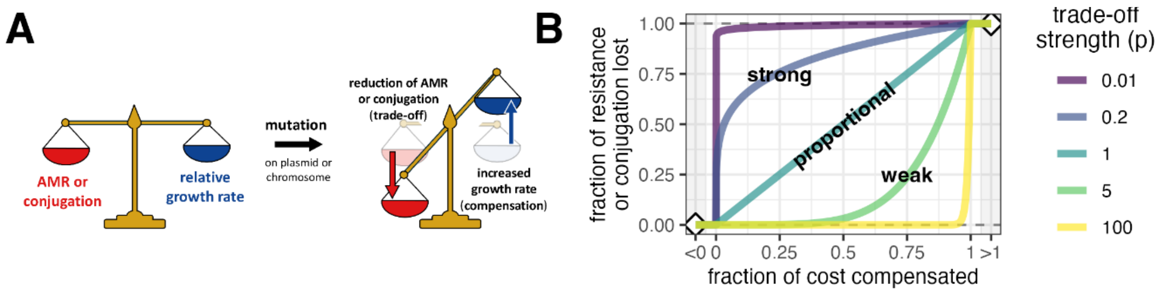


Figure S1. **Trade-offs associated with compensation between relative growth rate amelioration and reduction of conjugation or resistance. (A)** Mutations, irrespective of location, can cause increased relative growth rate by reducing plasmid-conferred antibiotic resistance or conjugation, which are metabolically costly. (**B**) The strength of a trade-off $p$ between fraction of cost compensated (x-axis) and fraction of resistance or conjugation lost (y-axis) can range from strong ($p = 0.01$, small compensation corresponds to big reduction) to proportional ($p = 1$) to weak ($p = 100$, only large compensation corresponds to big reduction). See Methods for details on the trade-off implementation.

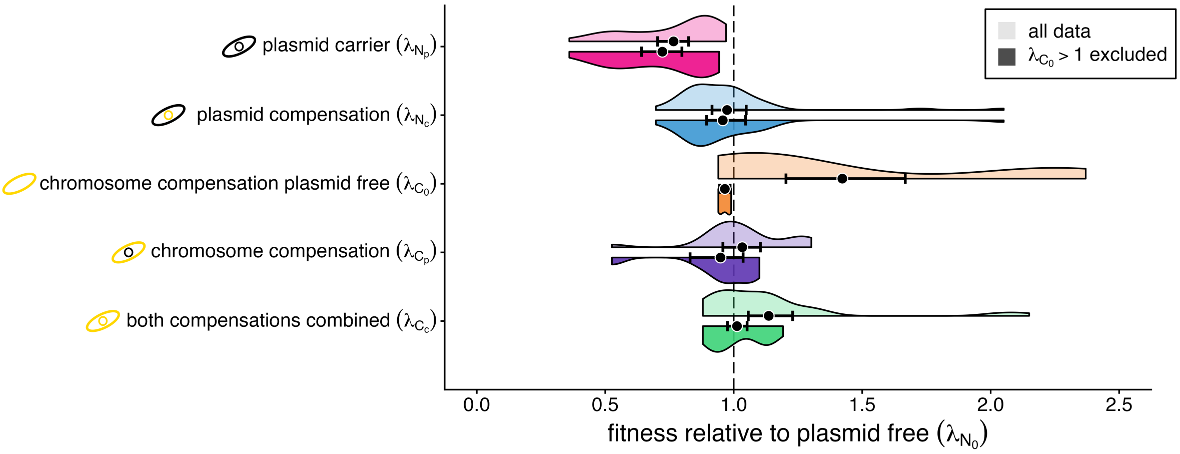


Figure S2. **Fitness costs and amelioration of plasmid carriage before and after experimental evolution, restricted to studies reporting Malthusian ratios from direct competition assays.** The lighter, upper halves of the violin plots show the kernel density estimates when all empirical data points are included and the darker, lower halves when cases where $C_0 > 1$ are excluded. Black points show means with bootstrapped 95% confidence intervals. See Figure 2 for detailed description. Raw data are in Supplemental Data 1 and summary statistics in Table S3.

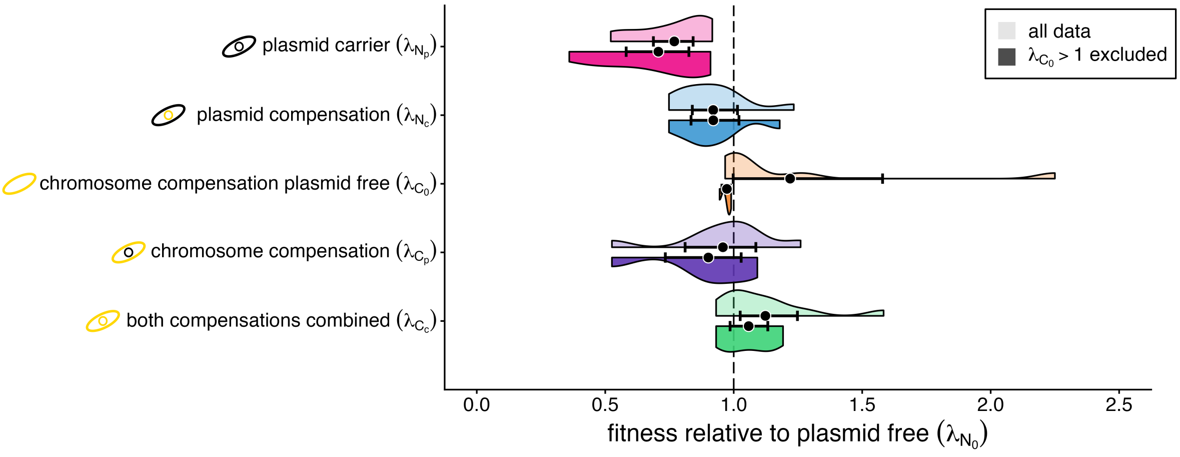


Figure S3. **Fitness costs and amelioration of plasmid carriage before and after experimental evolution, when averaged per study.** Shown are kernel density plots of mean fitness measures– one average value per study, across reported evolved lines or strains, rather than individual measurements (Figure 2, Figure S2). The qualitative finding of main Figure 2 is unchanged: $C_c$ displays the highest fitness, followed by $N_c$, then $C_p$. The lighter, upper halves of the violin plots show the kernel density estimates when all empirical data points are included and the darker, lower halves when cases where $C_0 > 1$ are excluded. Black points show means with bootstrapped 95% confidence intervals. See caption of Figure 2 for a detailed description. Raw data are in Supplemental Data 1 and summary statistics in Table S4.

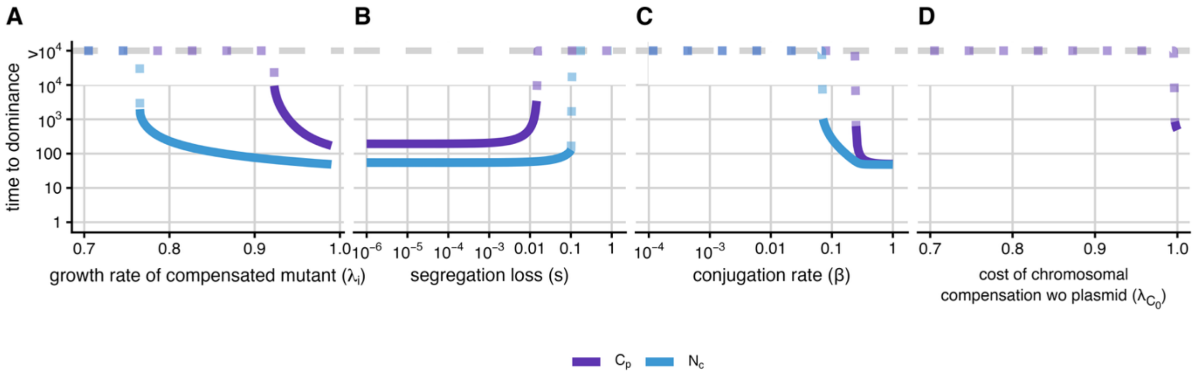


Figure S4. **In isolation, chromosomal compensation needs higher compensatory benefits than plasmid-borne compensation and is more limited by segregation loss, conjugation rate, and cost of chromosomal compensation.** Shown are the effects of different parameter values of (**A**) fitness associated with chromosome or plasmid compensation ($\lambda_i$), (B) segregation loss ($s$), (**C**) conjugation rate ($\beta$) and (**D**) the cost of chromosomal compensation in absence of plasmid ($\lambda_{C_0}$) on the time for compensated mutants to reach 0.5 population size (i.e., become dominant) for either plasmid-borne ($N_c$, blue) only or chromosomal compensation ($C_p$, purple) only. Starting from the default parameter sets used for simulations shown in Figure 4, we systematically varied one parameter at a time (either $s$, $\beta$, or $\lambda_{C_0}$). We did this for two isolated scenarios: either only plasmid-borne ($\mu_p = 0.000015, \mu_c = 0$) or only chromosomal compensatory mutations can emerge ($\mu_p = 0, \mu_c = 0.000015$). Note that benefits of compensatory mutation for chromosomal and plasmid compensation are identical ($\lambda_{C_p} = \lambda_{N_c} = 0.95$). For simplicity, in (**D**) only $C_p$ is shown because $\lambda_{C_0}$ has no effect on $N_c$ here. If, for a given parameter value, the compensated mutant did not reach a population size of ≥ 0.95 by the end of simulation (t=$10^4$ h) this is indicated by a dashed line extending to the "$>10^4$" label.

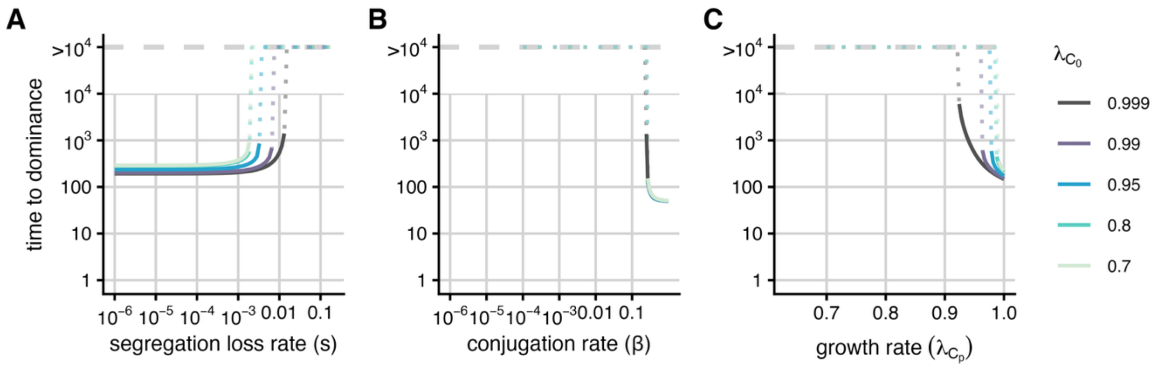


Figure S5. **Higher costs associated with chromosomal compensation in absence of the plasmid make chromosomal compensation more sensitive to changes in segregation loss, conjugation rate and compensatory benefit.** Starting from the default parameter set used for Figure 4C, we run simulations for varying values of: (**A**) segregation loss (s), (**B**) conjugation ($\beta$), and (**C**) growth rate of compensated mutants ($\lambda_{C_p}$). For each value of $s$, $\beta$ or $\lambda_{C_p}$ (x-axis), we plot the resulting time for chromosomally compensated mutants ($C_p$) to reach dominance, i.e., population size ≥ 0.5 (y-axis). Differently coloured lines indicate different values for the cost of chromosomal compensation in the absence of plasmid ($\lambda_{C_0}$) – (going from dark blue for low costs to light blue for high costs). If, for a given parameter value, $C_p$ did not reach a population size of ≥ 0.5 by t=$10^4$ h this is indicated by the line turning dashed and extending to the “>$10^4$” label.

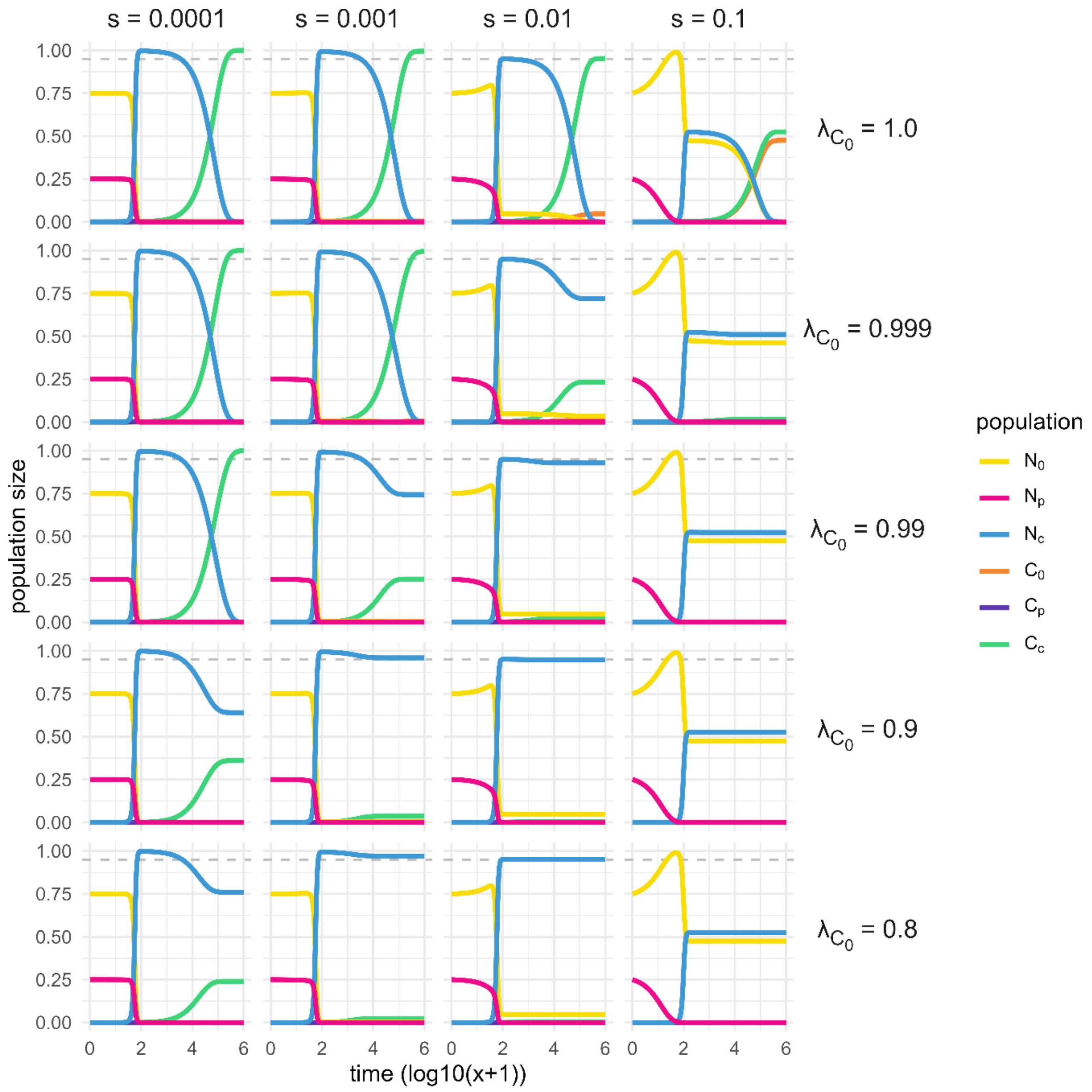


Figure S6. **Establishment and final frequency of cells with both compensatory mutations ($C_c$) depends on both segregation loss (s) and the cost of chromosomal compensation in absence of plasmid ($\lambda_{C_0}$).** Using the parameter set of Figure 4D as a baseline (both compensatory mutations are possible, and all compensatory mutations bring the same benefit), we vary s and $\lambda_{C_0}$ across selected values and plot the resulting dynamics for each parameter combination. The population sizes over time in h (log10(x+1)) are shown for: ancestral, plasmid-free cells ($N_0$, yellow), cells carrying the uncompensated plasmid ($N_p$, pink), cells carrying the compensated plasmid ($N_c$, blue), plasmid-free cells with a compensatory mutation on the chromosome ($C_0$, orange), and cells carrying the compensated plasmid as well as a compensatory mutation on the chromosome ($C_c$, green). A population size of 0.95 is indicated by the grey, dashed line.

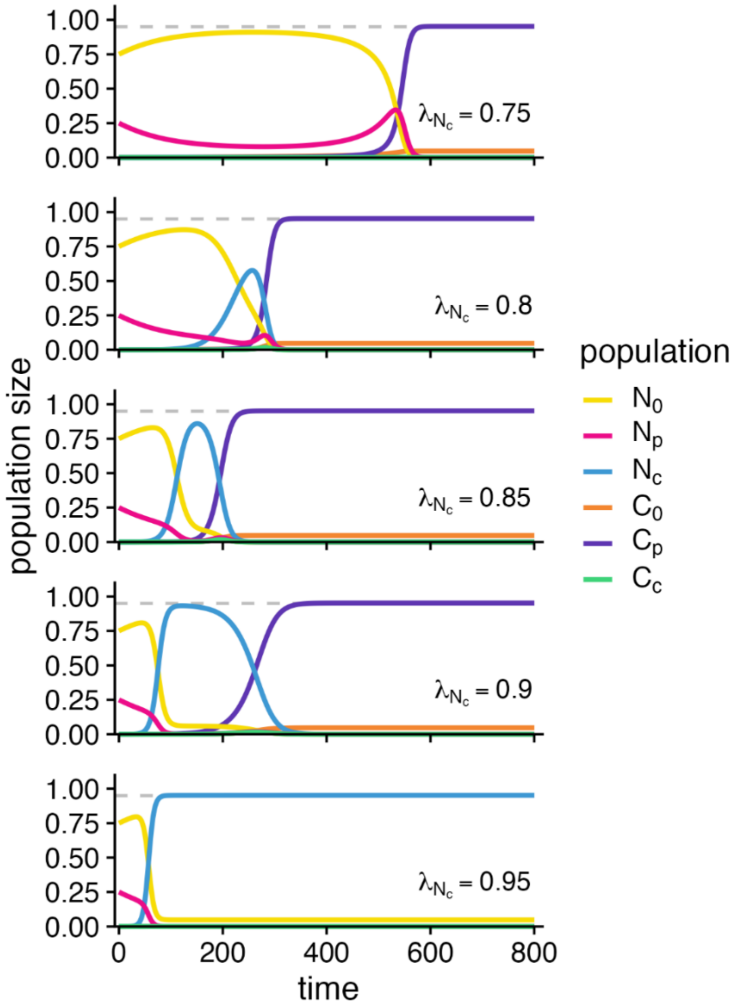


Figure S7. **The benefit of mobile compensation affects how quickly chromosomal compensation establishes in a non-linear manner.** Shown are simulations of compensatory evolution in which both chromosomal and plasmid-borne compensation are possible ($\mu_c = 1.5 \times 10^{-5}$, $\mu_p = 1.5 \times 10^{-5}$). Across panels, $\lambda_{N_c}$ increases from a value that provides no benefit relative to the ancestral plasmid-carrying population ($N_p$, pink) to a value equal to the benefit of chromosomal compensation ($C_p$), whose growth rate is fixed at $\lambda_{C_p} = 0.95$ (top to bottom: $\lambda_{N_c} = 0.75$, 0.8, 0.85, 0.9, 0.95). Depending on the value of $\lambda_{N_c}$ relative to $\lambda_{C_p}$ the presence of plasmid-borne compensation ($N_c$, blue) can either speed up or hinder the establishment of a population with chromosomal compensation ($C_p$, purple). All simulations start with $N_0 = 0.75, N_p = 0.25, N_c = C_0 = C_p = C_c = 0$. To isolate the comparison between $N_c$ and $C_p$, cells carrying both compensatory mutations simultaneously ($C_c$) are assumed to receive no compensatory benefit at all ($\lambda_{C_c} = \lambda_{N_p} = 0.75$). See Table S1 for all other parameters values. The grey line marks a population size of 0.95.

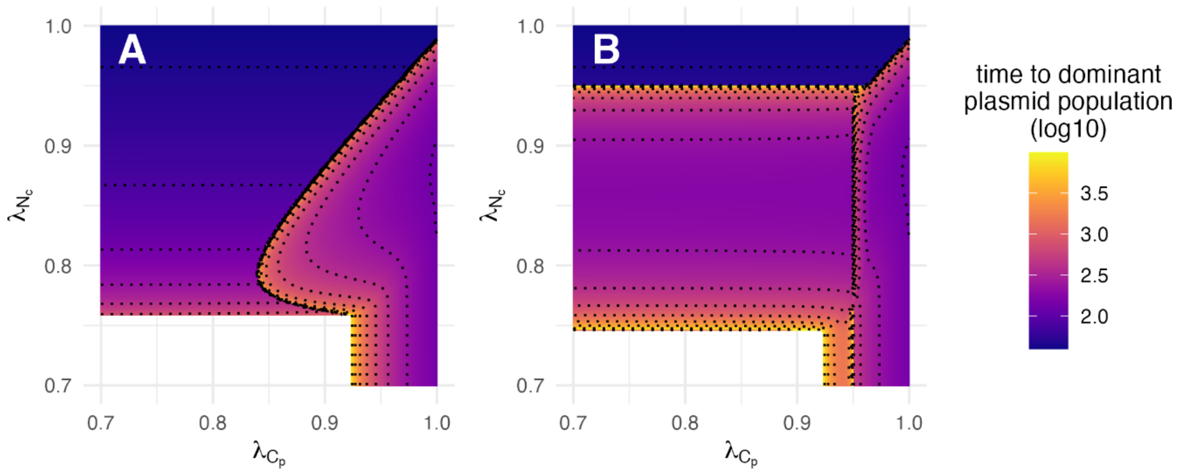


Figure S8. **The benefit of plasmid-borne compensation affects the time to dominance of chromosomal compensation, but not vice versa.** Simulation results across combinations of growth rates of chromosomal compensation $\lambda_{C_p}$ (x-axis) and of plasmid compensation $\lambda_{N_c}$ (y-axis) – each ranging from 0.7 to 1. For each parameter combination, the time to dominant (i.e., largest) plasmid population is shown on a log10 scale, which is the time point at which the plasmid-carrying population that remains dominant until the end of the simulation becomes dominant ($t_{end}$ = 10,000h). (**A**) Shows the scenario that the combination of both compensatory mutations simultaneously ($C_c$) provides no benefit ($\lambda_{N_p} = \lambda_{C_c} = 0.75$). (**B**) Shows the scenario where both compensatory mutations simultaneously ($C_c$) provides a high benefit ($\lambda_{C_c} = 0.95$). Together, this shows that time to dominance depends asymmetrically on the relative hierarchy of $\lambda_{N_c}$, $\lambda_{C_p}$, and $\lambda_{C_c}$ with $\lambda_{N_c}$ having the strongest effect. Intermediate values of $\lambda_{N_c}$ shorten time to dominance, whereas values close to $\lambda_{C_p}$ or $\lambda_{C_c}$ prolong it. All other parameters are kept constant (Table S1). The simulations are the same as shown in Figure 5. Note, that plasmid loss corresponds to the absence of a time value. Dashed contour lines show parameter combinations of $\lambda_{C_p}$ and $\lambda_{N_c}$ that result in equal time values.

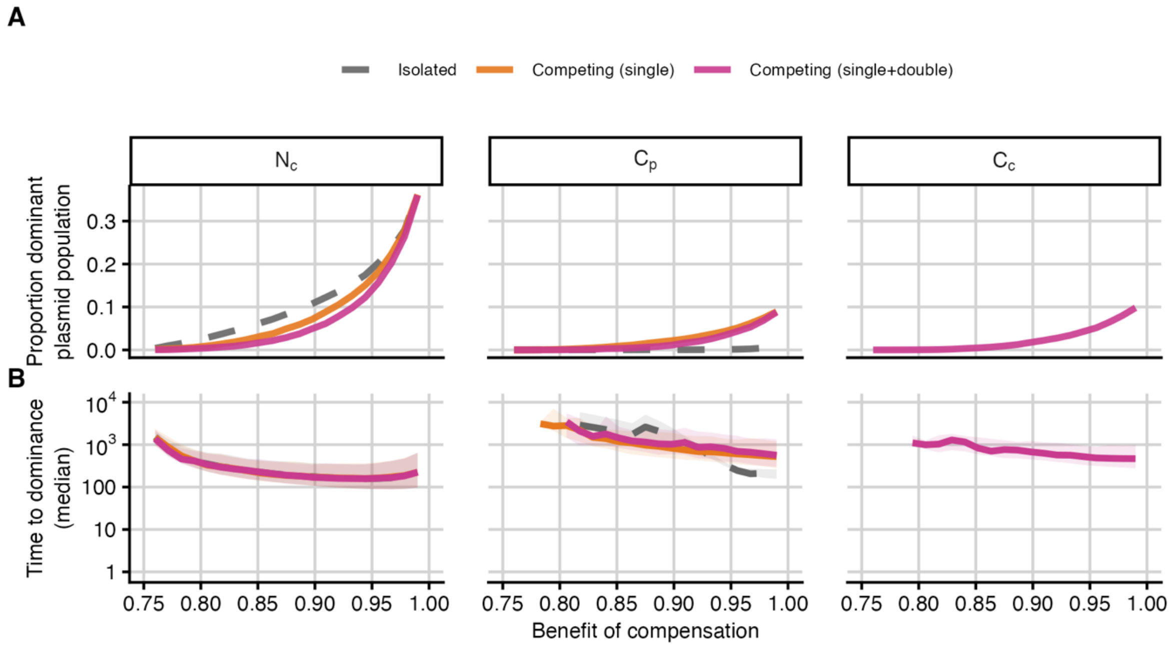


Figure S9. **Sensitivity analysis of time to dominance with respect to compensatory benefits and interactions between compensatory types across sampled parameter space.** Using Latin Hypercube Sampling we sampled 10,000 parameter sets from the ranges described in Table S6 – except for fixing growth rate of the ancestral plasmid-carrying bacteria ($\lambda_{N_p}$ = 0.75). For each parameter set, we ran simulations while systematically increasing the growth rate (thereby the benefit of compensation) of only the focal compensatory type – $\lambda_{N_c}$, $\lambda_{C_p}$, or $\lambda_{C_c}$ – from 0.76 to 0.99 in 21 steps. We did this for three scenarios: "Isolated", "Competing (single)", "Competing (single+double)". In "Isolated", only chromosomal or plasmid-borne compensation can arise (by setting either $\mu_p$ or $\mu_c$ set to zero, respectively). In "Competing (single)", $N_c$ and $C_p$ can both arise, but $C_c$ carries no fitness benefit ($\mu_c = \mu_c = 1.5 \times 10^{-5}$ and $\lambda_{C_c} = \lambda_{N_p} =$ 0.75) and hence cannot establish. In "Competing (all)", $\lambda_{C_c}$ is also sampled from 0.76–0.99, and hence all three compensated mutants compete and can establish. (A) Proportion of simulations in which the focal compensatory type is the dominant plasmid-carrying population at the end of the simulation ($t_{end}$ = 10,000). (B) Median time to dominance of the focal population. The shaded areas show interquartile ranges.

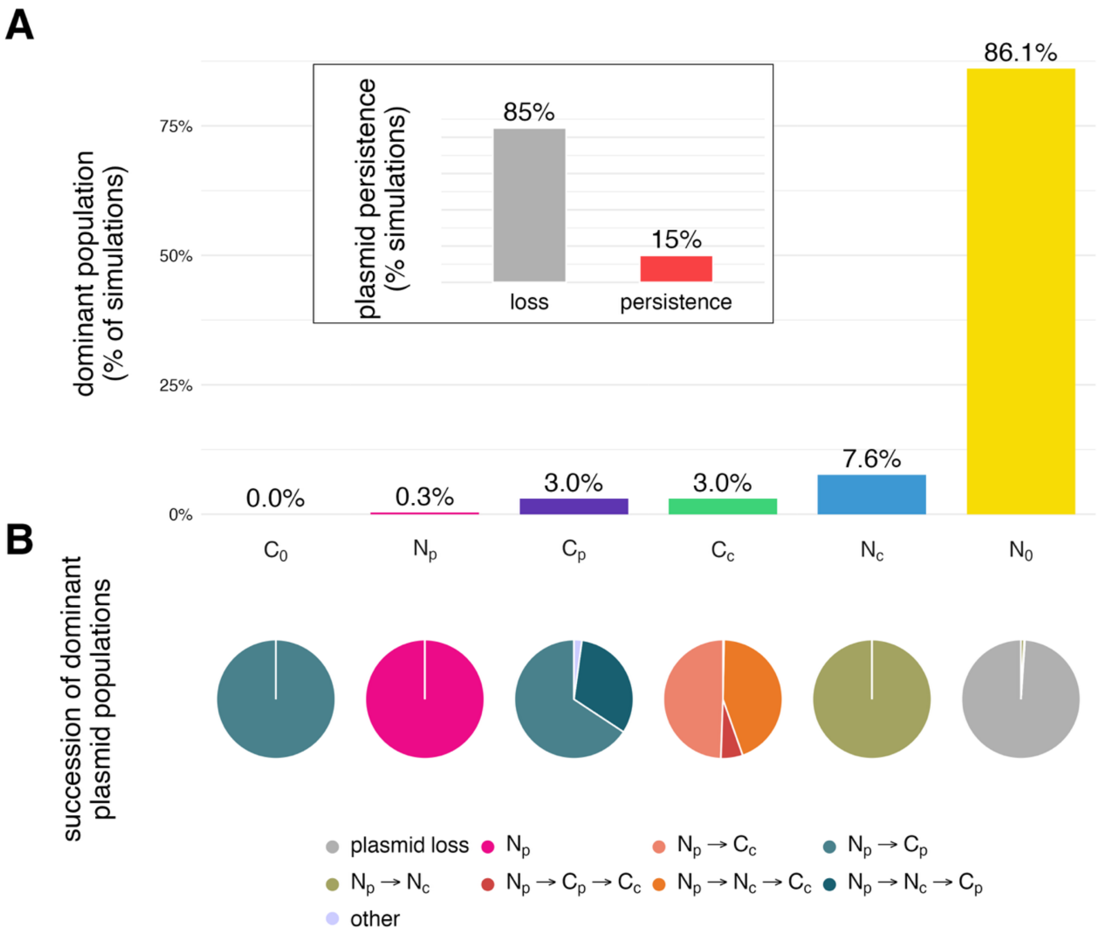


Figure S10. **Conditional uniform sampling: Plasmid compensation most often drives plasmid persistence and is typically transiently dominant before chromosomal or both compensations.** Shown are simulation results of 100,000 parameter sets, each including at least one compensated mutant with higher growth rate than the ancestral plasmid-carrying population $N_p$ (see Text S2 for details). (**A**) Percentage of simulations in which the plasmid-free ($N_0$), uncompensated plasmid ($N_p$), plasmid compensation ($N_c$), chromosomal compensation without plasmid ($C_0$), chromosomal compensation ($C_p$), or both compensations ($C_c$) populations are dominant – i.e., the largest – at the end of simulation ($t_{end}$ = 10,000 h). (**B**) Successions leading to final dominant populations. For each final dominant population in A, the corresponding pie chart shows which, if any, plasmid populations succeeded each other as dominant plasmid population. Plasmid loss is defined as the total size of plasmid-carrying populations at $t_{end}$ being less than 0.001. See inset for overall percentages of simulations that resulted in plasmid loss or persistence. Successions occurring in less than 1% of corresponding simulations are grouped as "other".

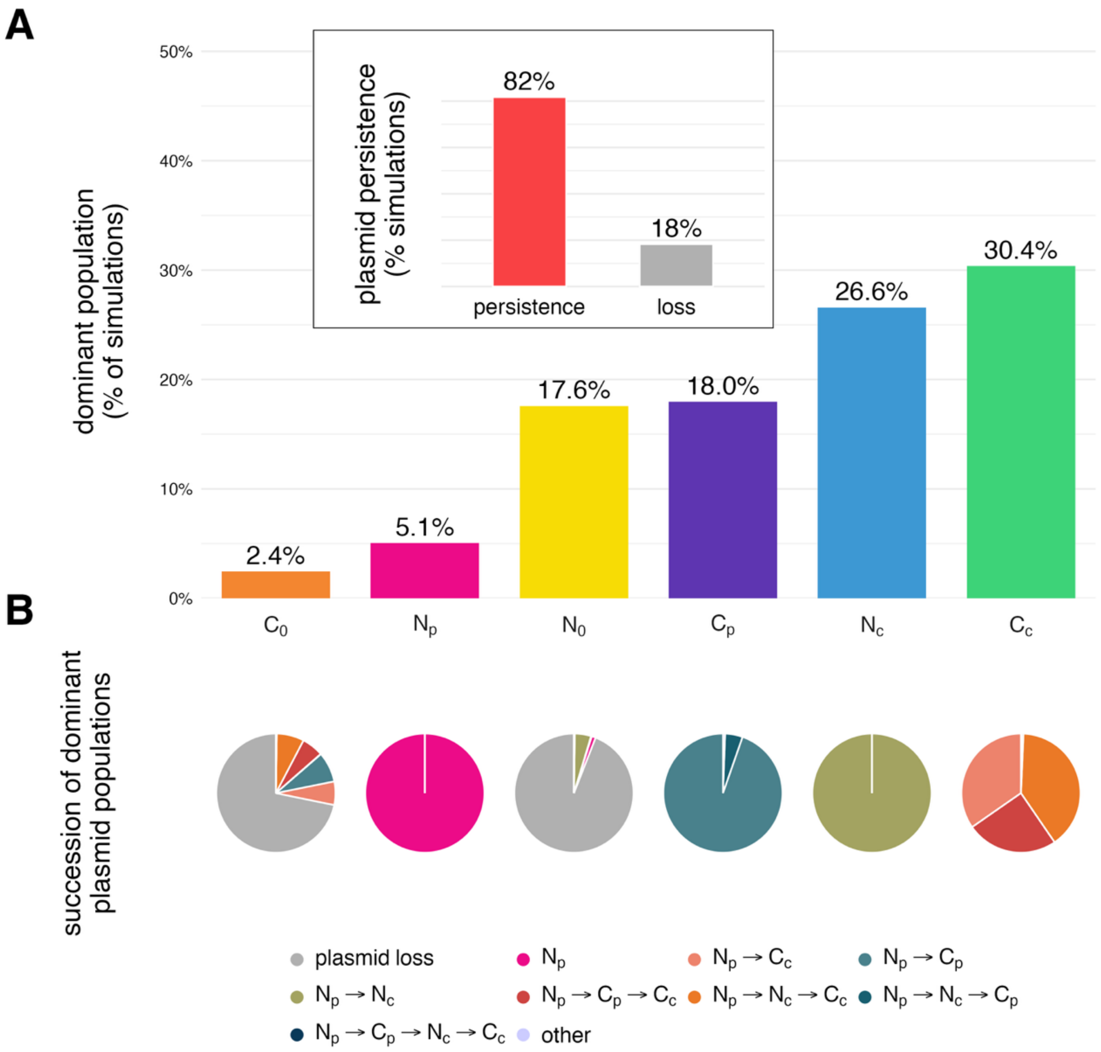


Figure S11. **Gamma Sampling: Growth rates from empirical distributions increase plasmid persistence via combined compensation, while plasmid compensation remains the most frequent dominant plasmid bearing population when transient and ultimate dominance are summed.** Shown are simulation results of 100,000 parameter sets sampled from gamma distributions fitted to the empirical data (Figure S2). See Text S2 for a detailed description of the sampling scheme. Shapes and rates of gamma distribution can be found in Table S6. (**A**) Percentage of simulations in which $N_0$, $N_p$, $N_c$, $C_0$, $C_p$, or $C_c$ are the dominant populations at the end of the simulation ($t_{end}$ = 10,000 h). (**B**) Successions leading to the corresponding final dominant populations. See Figure S10 for detailed panel descriptions.

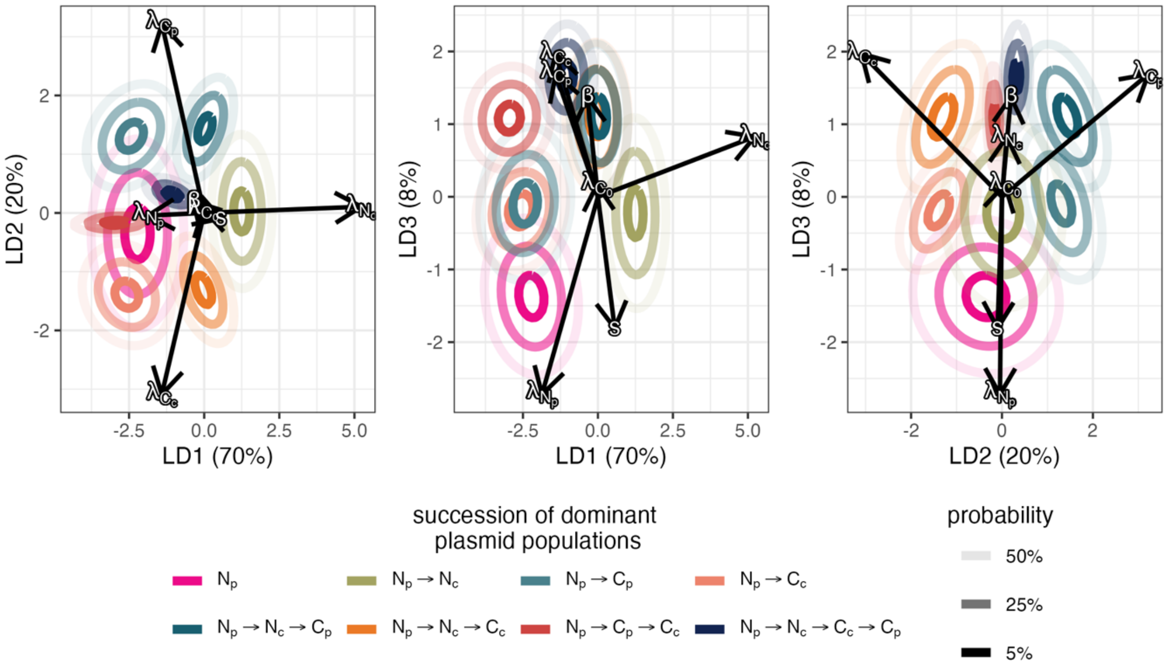


Figure S12. **Conditional uniform sampling: Sensitivity of successions to model parameters using Linear Discriminant Analysis (LDA).** LDA of the successions of dominant plasmid populations from 100,000 simulations (same simulation results as shown in Figure S10 but filtering out plasmid loss). Arrow length indicates the importance and arrow angle the direction of model parameters in class separation (see Table S1 for parameter descriptions). Contours show the x% highest-density regions of a bivariate normal fitted per succession, drawn as Gaussian ellipses for visual clarity. See Text S2 for details on LDA and Table S6 for ranges of parameter sampling. Growth rates $\lambda_i$ (reflecting plasmid cost and compensatory benefits) have the strongest influence showing that successions are driven by the hierarchy of growth rates.

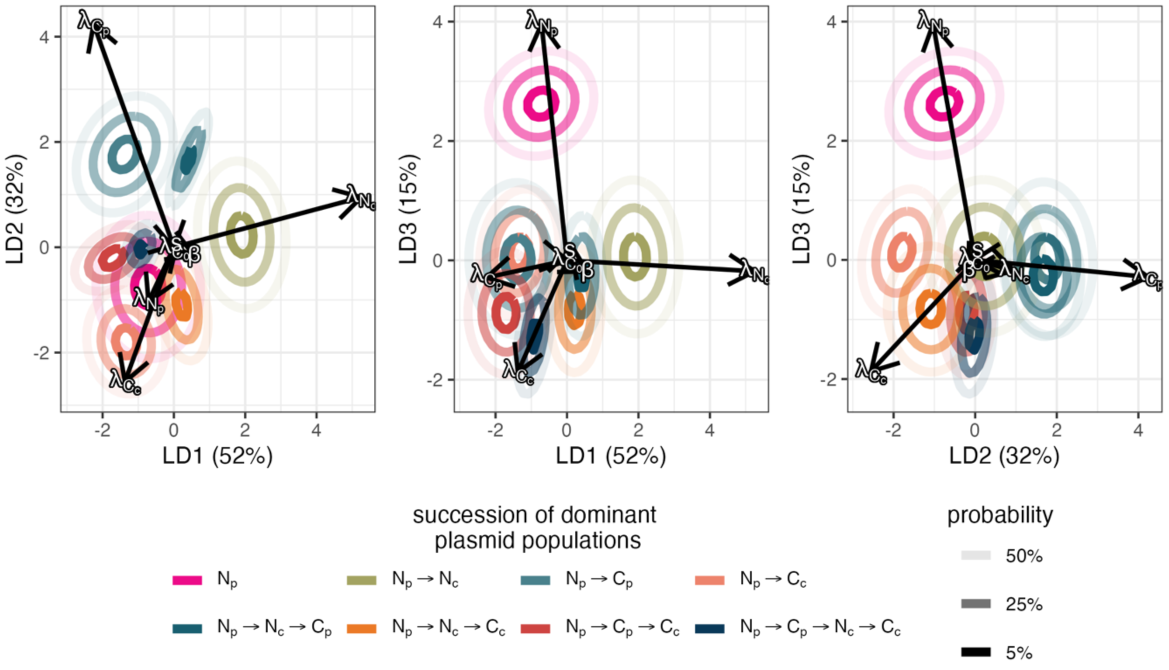


Figure S13. **Gamma sampling: Sensitivity of successions to model parameters using Linear Discriminant Analysis (LDA).** LDA of the successions of dominant plasmid populations from 100,000 simulations (same simulation results as shown in Figure S11 but filtering out simulations where plasmid loss occurred). The length of arrow indicates the importance and arrow angle the direction of the corresponding model parameters in class separation (see Table S1 for parameter descriptions). Contours show the x% highest-density regions of a bivariate normal fitted per succession, drawn as Gaussian ellipses for visual clarity. See Text S2 for details on LDA and Table S6 for distributions of gamma sampling. Growth rates $\lambda_i$ retain the strongest influence in separating the successions, whether sampled from empirical or uniform distributions (Figure S12).

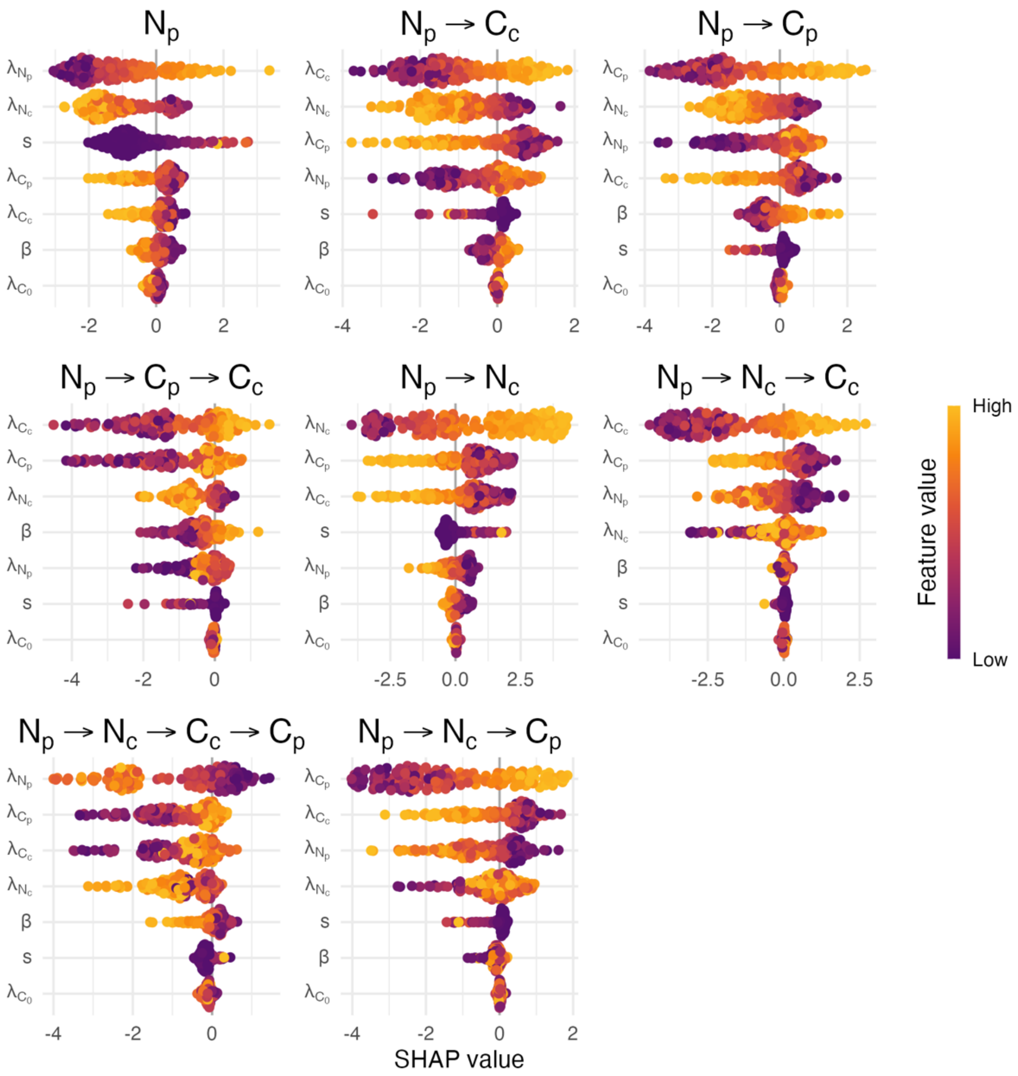


Figure S14. **Conditional uniform sampling: Sensitivity of successions to model parameters using gradient-boosted decision trees (XGBoost) and Shapley Additive Explanatory (SHAP) values.** Beeswarm plots showing SHAP values for an XGBoost classifier trained on the successions of dominant plasmid populations from 100,000 simulations (same simulations as in Figure S10 but filtering out plasmid loss). Each subplot shows the parameter importance (ranked by mean |SHAP| value, most important at top) and direction for the depicted succession. Positive SHAP values reflect that the parameter increases the predicted probability of that succession and negative values reflect a decrease. The colour of the points indicates the feature value, i.e., the model parameter value, from low (purple) to high (yellow). See Table S1 for descriptions, Table S6 for distributions of model parameters, and Text S2 for details on implementation of XGBoost and SHAP values. Growth rates $\lambda_i$ consistently rank as the strongest SHAP features, highlighting the importance of growth rate hierarchy in determining which succession occurs.

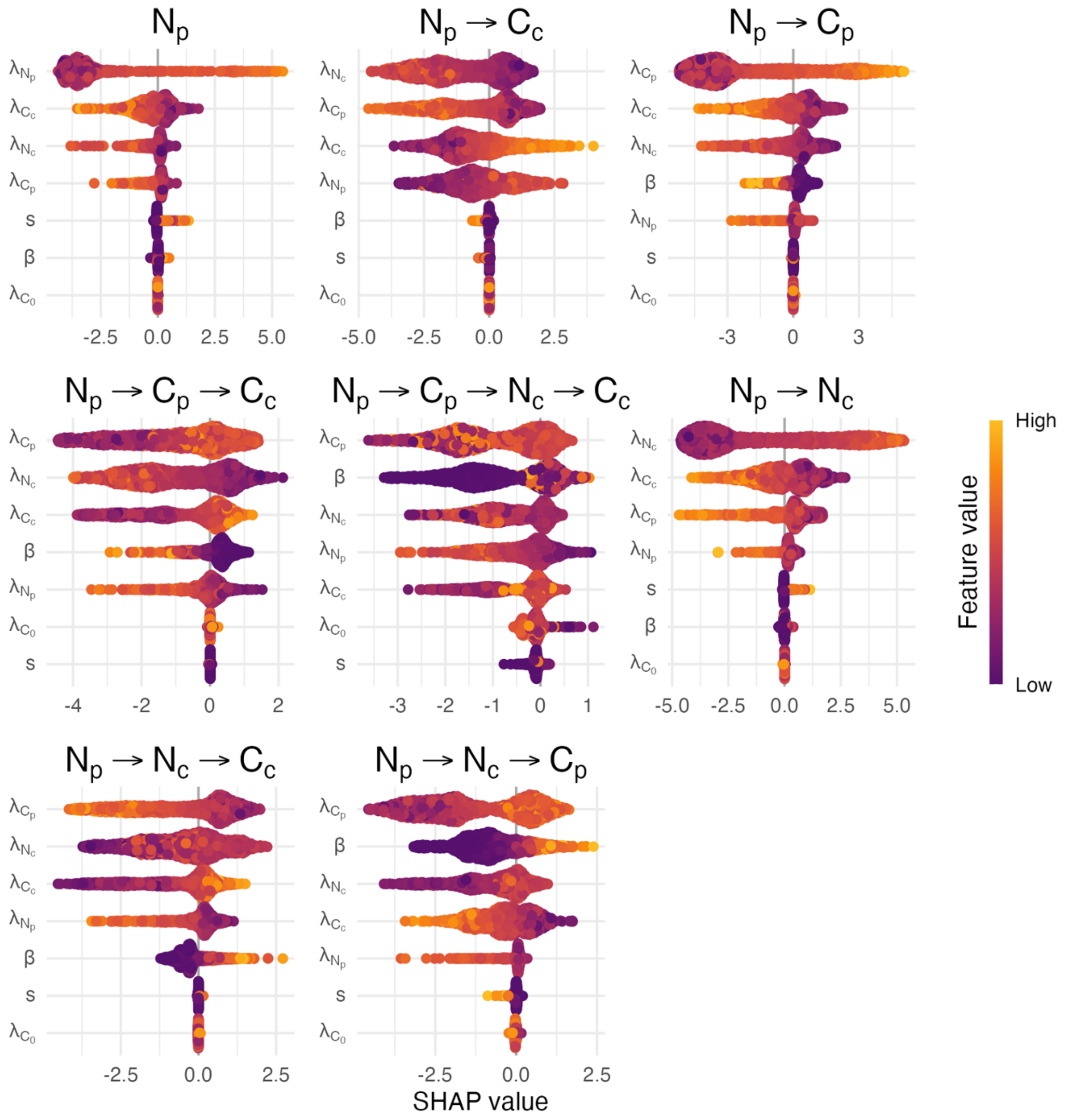


Figure S15. **Gamma sampling: Sensitivity of successions to model parameters using XGBoost and SHAP values.** Beeswarm plots based on the same 100,000 simulations shown in Figure S11 (excluding plasmid loss). Features represent model parameters ranked by mean |SHAP| value (most important at the top). The sign of the SHAP value indicates whether the parameter increases or decreases the probability of predicting the corresponding succession, and point colours reflect relative parameter values (low (purple) to high (yellow)). Mirroring the results from conditional uniform sampling (Figure S14), growth rates $\lambda_i$ consistently rank as the top SHAP features, showcasing their central role in determining succession outcomes. See Figure S14 and Text S2 for a more detailed description of XGBoost and SHAP values.

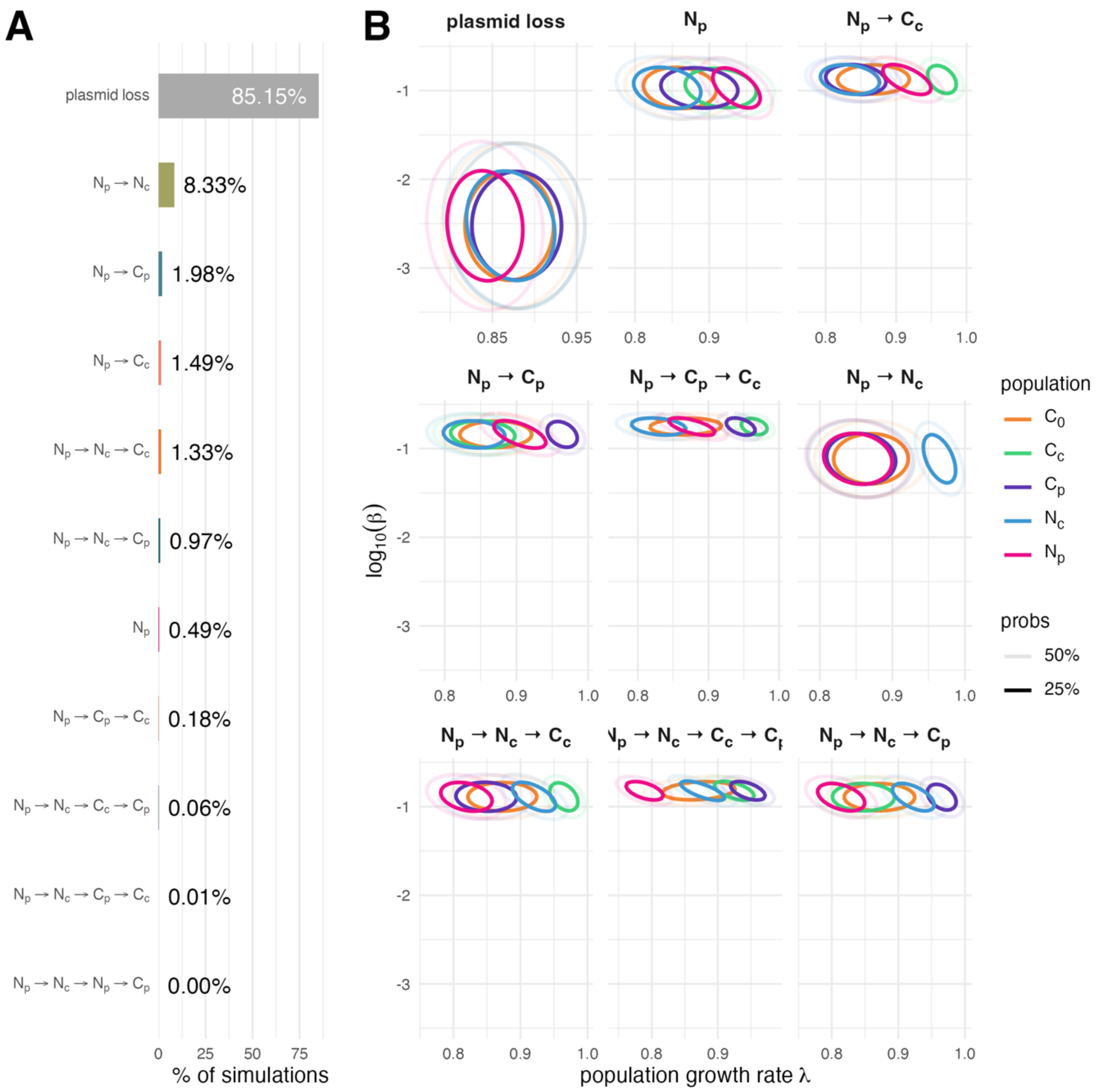


Figure S16. **Growth rate $\lambda$ and conjugation rate $\log_{10}(\beta)$ distributions per population by succession of dominant plasmid populations.** (**A**) Percentages of simulations by succession (100,000 simulations with sampled parameter sets same simulations as in Figure S10). (**B**) For each succession (excluding successions below 0.1%), density contours approximate where each populations growth rate ($\lambda$, x-axis) and conjugation rate (β, y-axis) values fall across simulations. The contours enclose the highest density regions containing 25% and 50% of the estimated probability mass. This illustrates that plasmid loss is primarily determined by conjugation rate β, and which succession occurs is then determined by the hierarchy of growth rates $\lambda_i$, with higher growth rates occurring later in the succession.

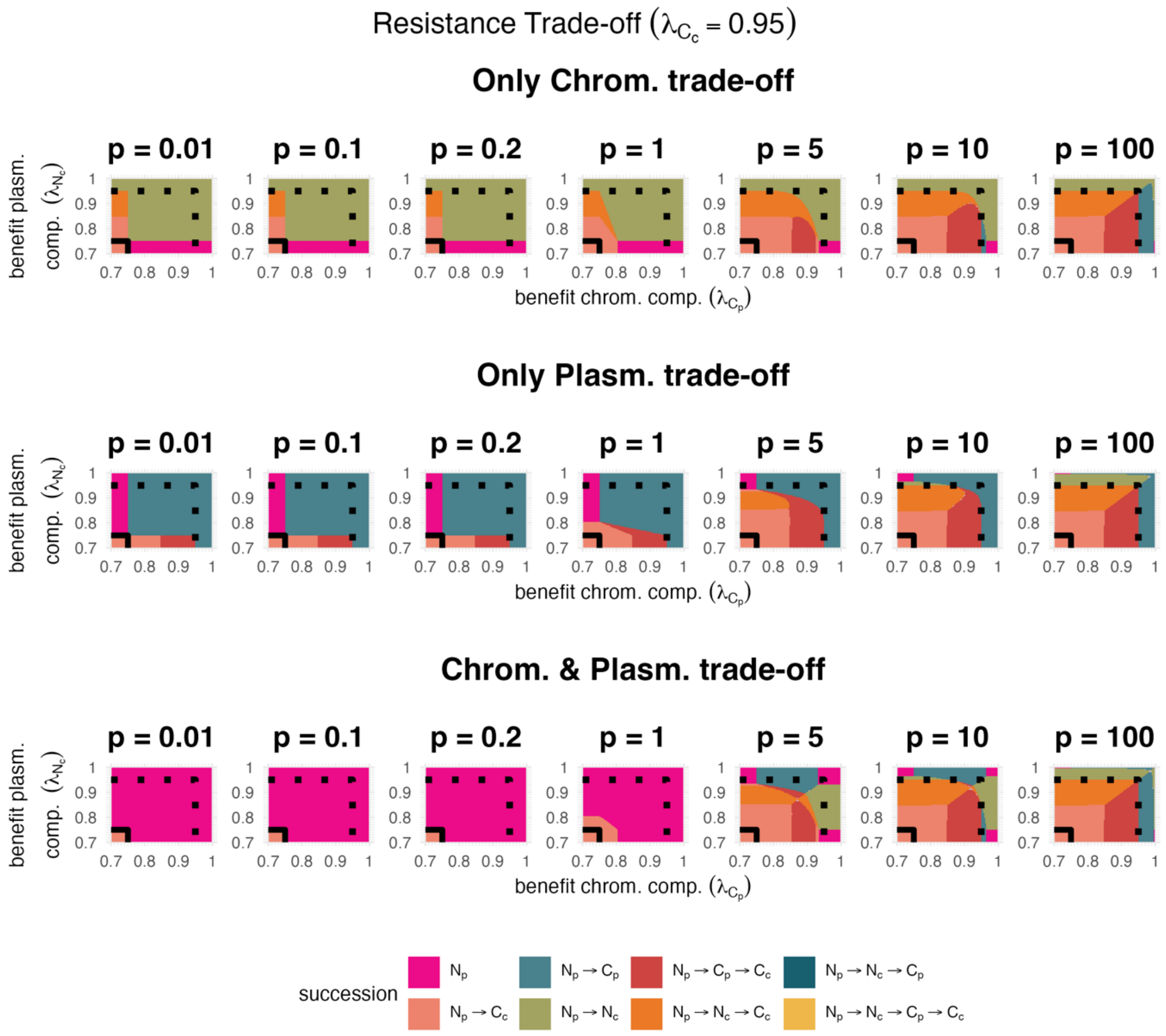


Figure S17. **Resistance trade-offs across trade-off strengths, for the scenario that both compensatory mutations simultaneously confer a high benefit ($\lambda_{C_c} = 0.95$).** Individual heatmaps show the succession of dominant plasmid populations across combinations of relative growth rates of cells with compensatory mutations on the plasmid $\lambda_{N_c}$ (y-axis) and on the chromosome $\lambda_{C_p}$ (x-axis). See Figure 5 for details on the successions. Shown are three different scenarios as in Figure 6: the trade-off is associated with mutations on the chromosome only (top row), the plasmid only (middle row), or both (bottom row) – each for increasing trade-off strengths from sub-proportional ($p = 0.01$) to super-proportional ($p = 100$). Trade-offs act on resistance under antibiotic selection ($A = 1$). See Methods for details on the implementation and Figure S1 for a visualisation of the trade-off function.

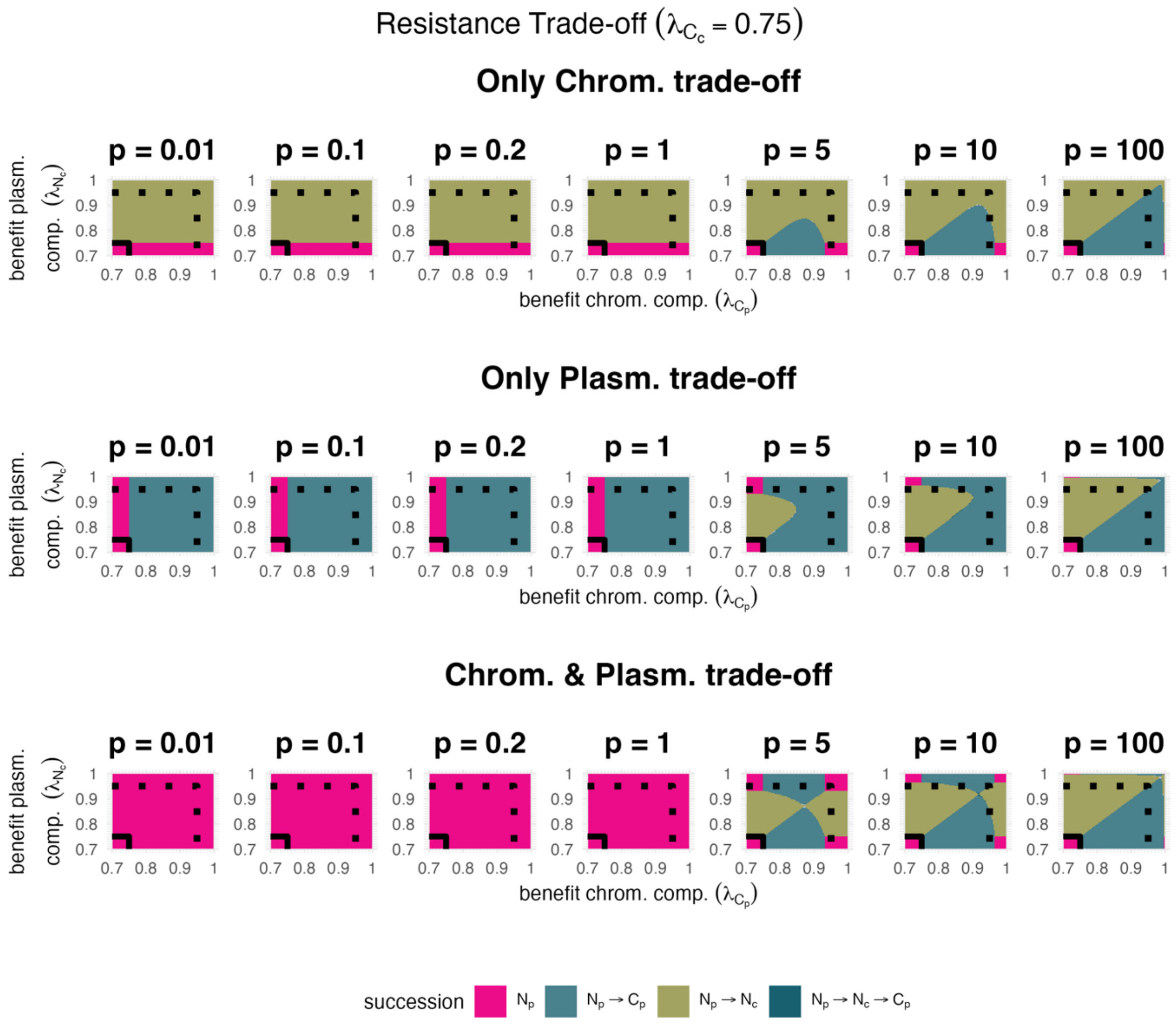


Figure S18. **Resistance trade-offs across trade-off strengths, for the scenario that both compensatory mutations simultaneously confer no benefit relative to the ancestral plasmid cost ($\lambda_{C_c} = 0.75$).** See Figure S17 for description.

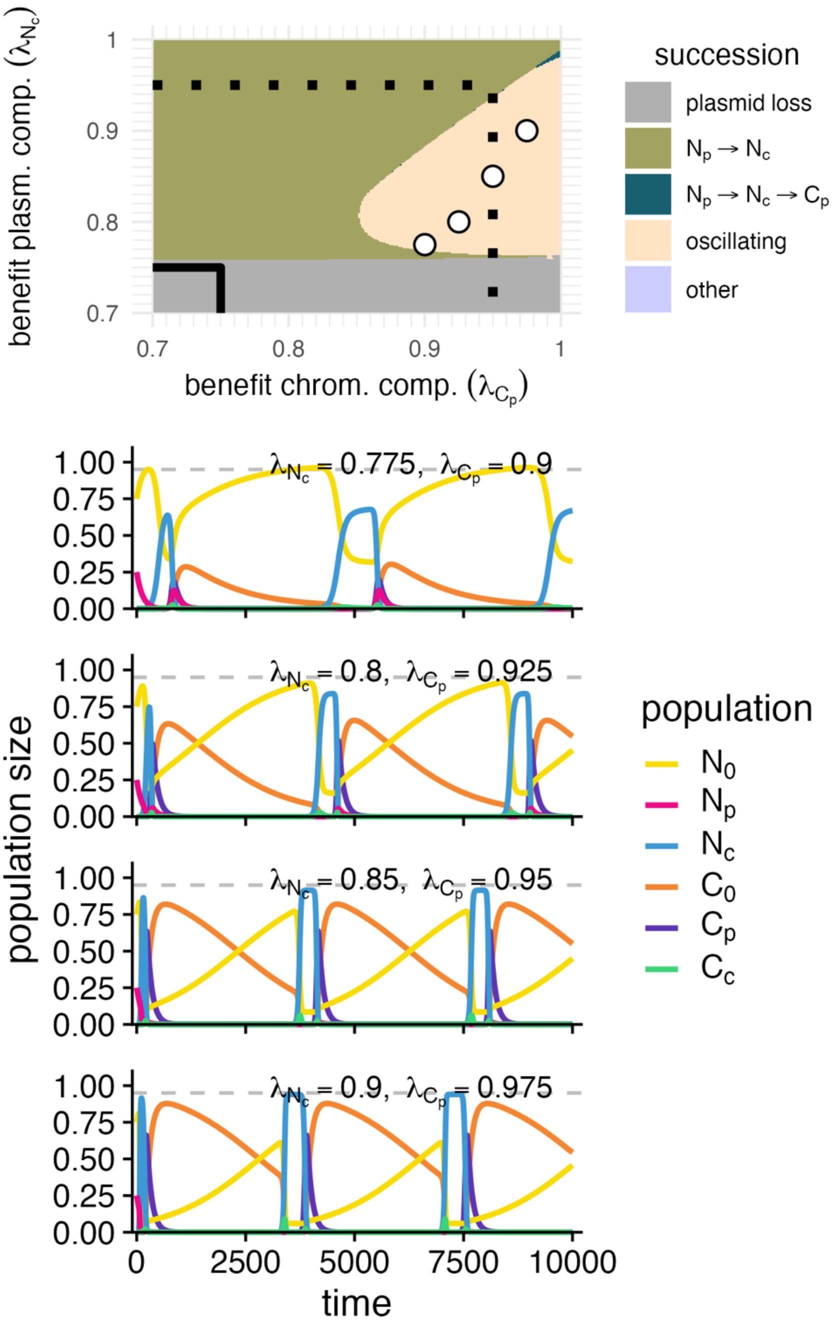


Figure S19. **Examples of oscillatory dynamics.** Oscillatory population dynamics can occur if conjugation rates between compensated mutants differ. Shown are simulations for selected pairs of $\lambda_{N_c}$ and $\lambda_{C_p}$ indicated by point in the heatmap on top, under a conjugation trade-off associated with chromosomal compensation (p=1, same as in Figure 6).

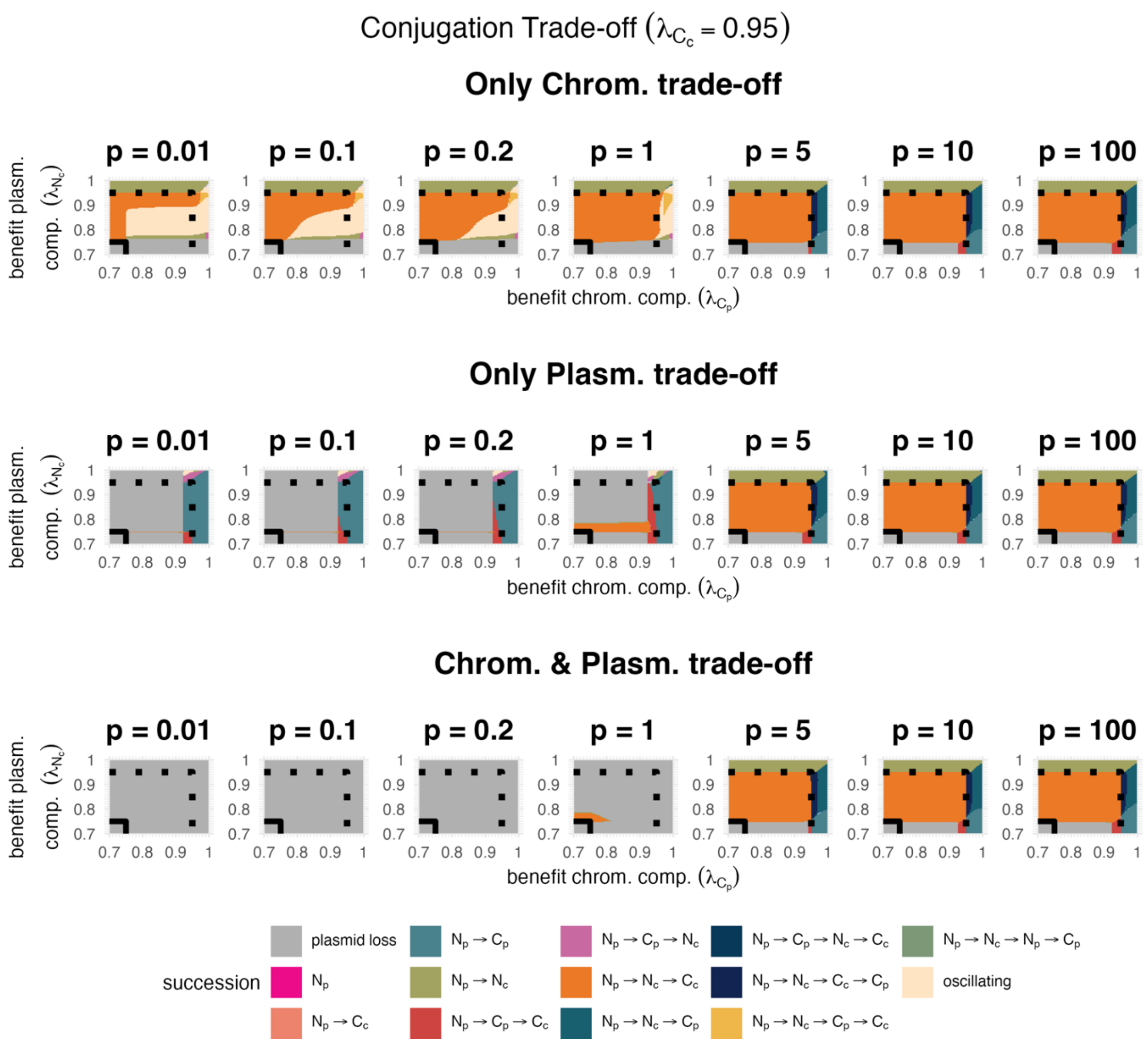


Figure S20. **Conjugation trade-offs across trade-off strengths, for the scenario that both compensatory mutations simultaneously confer a high benefit ($\lambda_{C_c} = 0.95$).** Conjugation trade-offs are investigated in absence of antibiotic selection ($A = 0$). Otherwise, see Figure S17 for description.

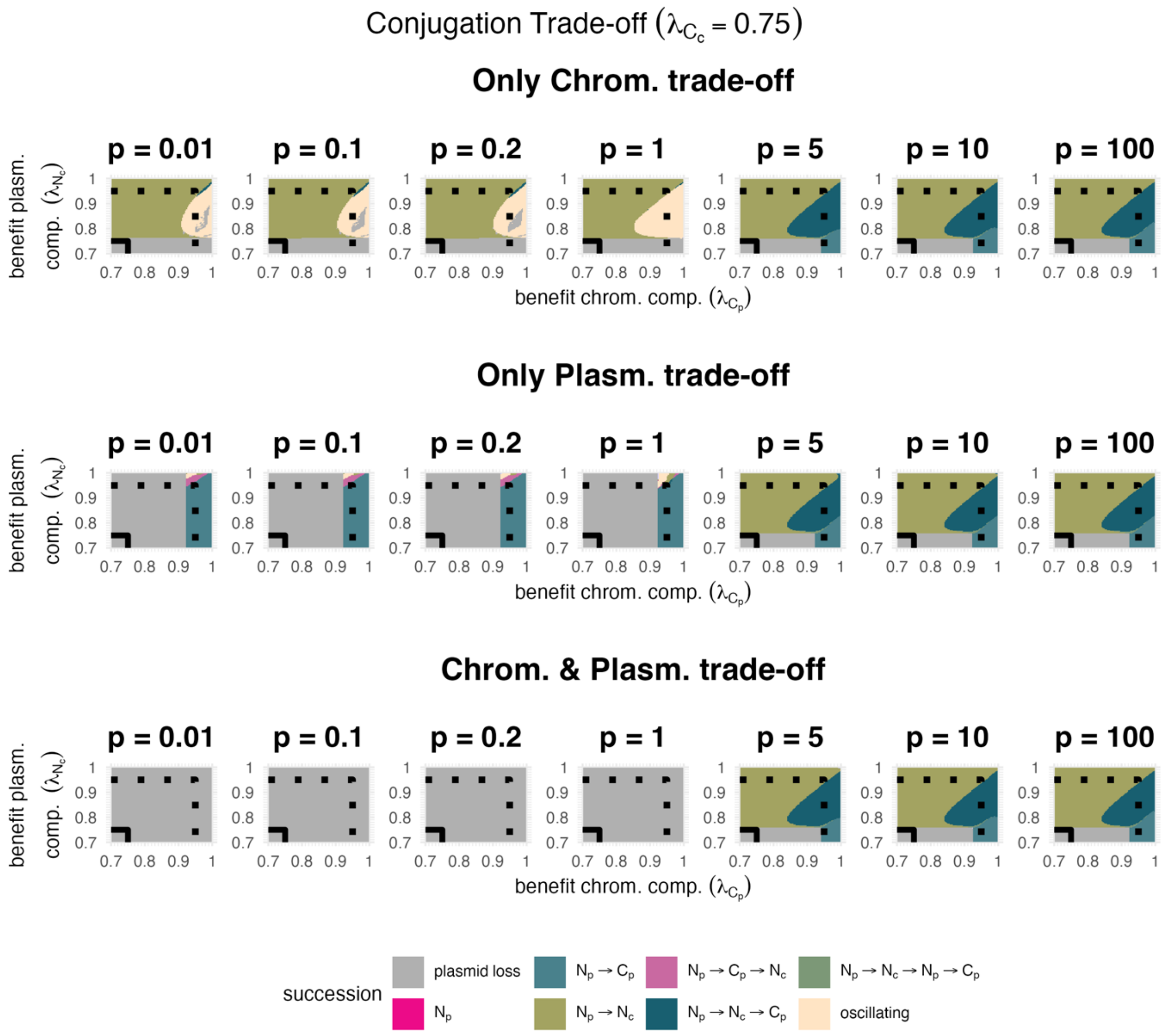


Figure S21. **Conjugation trade-offs across trade-off strengths, for the scenario that both compensatory mutations simultaneously confer no benefit relative to the ancestral plasmid cost ($\lambda_{C_c} = 0.75$).** Conjugation trade-offs are investigated in absence of antibiotic selection ($A = 0$). Otherwise, see Figure S17 for description.
